# Small molecule modulation of a redox-sensitive stress granule protein dissolves stress granules with beneficial outcomes for familial amyotrophic lateral sclerosis models

**DOI:** 10.1101/721001

**Authors:** Hiroyuki Uechi, Sindhuja Sridharan, Jik Nijssen, Jessica Bilstein, Juan M. Iglesias-Artola, Satoshi Kishigami, Virginia Casablancas-Antras, Ina Poser, Eduardo J. Martinez, Edgar Boczek, Michael Wagner, Nadine Tomschke, António M. de Jesus Domingues, Arun Pal, Thom Doeleman, Sukhleen Kour, Eric Nathaniel Anderson, Frank Stein, Hyun O. Lee, Xiaojie Zhang, Anatol W. Fritsch, Marcus Jahnel, Julius Fürsch, Anastasia C. Murthy, Simon Alberti, Marc Bickle, Nicolas L. Fawzi, André Nadler, Della C. David, Udai B. Pandey, Andreas Hermann, Florian Stengel, Benjamin G. Davis, Andrew J. Baldwin, Mikhail M. Savitski, Anthony A. Hyman, Richard J. Wheeler

## Abstract

Neurodegeneràve diseases such as amyotrophic lateral sclerosis (ALS) are often associated with mutàons in proteins that are associated with stress granules. Stress granules are condensates formed by liquid-liquid phase separàon which, when aberrant, can lead to altered condensàon behaviours and disease phenotypes. Here, we identified lipoamide, a small molecule which specifically prevents cytoplasmic condensàon of stress granule proteins. Thermal proteome profiling showed that lipoamide preferentially stabilises intrinsically disordered domain-containing proteins. These include SRSF1 and SFPQ, stress granule proteins necessary for lipoamide activity. The redox state of SFPQ correlates with its condensate-dissolving behaviour, in concordance with the importance of the dithiolane ring for lipoamide activity. In animals, lipoamide ameliorates aging-associated aggregàon of a stress granule reporter, improves neuronal morphology, and recovers motor defects caused by expression of ALS-associated FUS and TDP-43 mutants. In conclusion, lipoamide is a well-tolerated small molecule modulator of stress granule condensàon and dissection of its molecular mechanism identified a cellular pathway for redox regulàon of stress granule formàon.

## Introduction

Amyotrophic lateral sclerosis (ALS) is a fatal neurodegeneràve disease, primarily affecting motor neurons, with poor prognosis and few options for therapy^1^. Currently three FDA approved drugs are available: riluzole, edaravone, and, recently approved, relyvrio™ (a combinàon of sodium phenylbutyrate and taurursodiol)^2–4^. However, none blocks disease progression, and thus investigàng new therapeùc routes is important to overcome ALS. Many mutàons associated with familial ALS are found in RNA-binding proteins. Notably, TAR DNA-binding protein 43 (TDP-43) and fused in sarcoma (FUS), with >40 ALS-associated mutàons described in each^5–7^. These RNA-binding proteins have large intrinsically disordered regions (IDRs) with low sequence complexity.

TDP-43 and FUS are examples of stress granule proteins which normally localise to the nucleus, where they have crucial functions in gene expression regulàon and DNA damage responses. For example, FUS localises to paraspeckles and DNA damage foci in the nucleus^8,9^. Upon cellular stress, TDP-43 and FUS are exported to the cytoplasm where they become incorporated into stress granules, although neither are necessary for stress granule formàon^10^. Stress granules are liquid-like cytoplasmic assemblies, or condensates, which are formed by liquid-liquid phase separàon of both nuclear exported and constitùvely cytoplasmic proteins, along with mRNA^11–13^. Stress granule formàon is triggered by cellular stresses, such as oxidàve stress. This is often dependent on the cytoplasmic stress granule protein G3BP1^10^. When the cellular stress is alleviated, stress granules dissolve and proteins that normally reside in the nucleus, including FUS and TDP-43 translocate back to the nucleus.

It has been proposed that ALS-linked FUS and TDP-43 mutants cause diseases in part by inducing aberrant phase transìon of stress granules^12,14,15^. This reduces the dynamics of stress granule proteins, preventing them from dissolving when stress is removed, thus trapping nuclear proteins in the cytoplasm. Supportive evidence is that FUS and TDP-43 mutants often show constitùve mislocalisàon to the cytoplasm, and FUS tends to aggregate more readily in the cytoplasm^16^. Both mechanisms may cause a loss-of-function phenotype in the nucleus or a gain-of-function (cytotoxic) phenotype in the cytoplasm as cytoplasmic aggregates or fibrils: these are associated with motor neuron dysfunction leading to neurodegeneràve disease^17–19^. Either way, dissolving aberrant stress granules, reducing sensìvity to triggers of stress granule formàon, preventing or reversing stress granule protein aggregàon, and/or driving protein back to the nucleus might be an efficient way to prevent or reverse the consequences of ALS.

Small molecules have been an essential tool for relàng cytology to function. For instance, the microtubule polymerizàon inhibitor nocodazole have helped us to study cytokinesis and microtubule-mediated intracellular trafficking. Inhibitors against actin/myosin II activìes such as cytochalasin and Y-27632 have illuminated pivotal roles of these cytoskeletal proteins in cell shape, division, and migràon. Indeed compounds have been identified that can disrupt stress granule condensàon, especially 1,6-hexanediol^20^ and similar alcohols^21^. However, these compounds are both toxic and non-specific, as they affect multiple condensates^21,22^. There is an unmet need for non-toxic and specific compounds able to specifically dissolve condensates. Here we searched for compounds that both prevented the formàon of stress granules, and induced their dissolùon, and identify lipoamide, which partitions into stress granules in cells and alleviates pathology caused by ALS-associated FUS and TDP-43 mutants in both motor neurons *in vitro* and in fly models of ALS. Using lipoamide as a tool compound, we identify a pathway that allows stress granules to sense the oxidàve state of the cell.

## Results

We performed a cell-based screen of 1,600 small molecules to identify compounds which affect stress granule formàon upon arsenate treatment, by monitoring GFP-tagged FUS (FUS-GFP) localisàon in HeLa cells (Fig. 1A–C). Many compounds altered FUS-GFP localisàon in stressed cells, often reducing the number of FUS-GFP-containing stress granules (Fig. 1B,C). Emène, known to prevent stress granule formàon by stabilising polysomes^23^, was present in the library and acted as a posìve control. Edaravone, an FDA-approved ALS therapeùc^24^, had no significant effect.

**Fig. 1.**
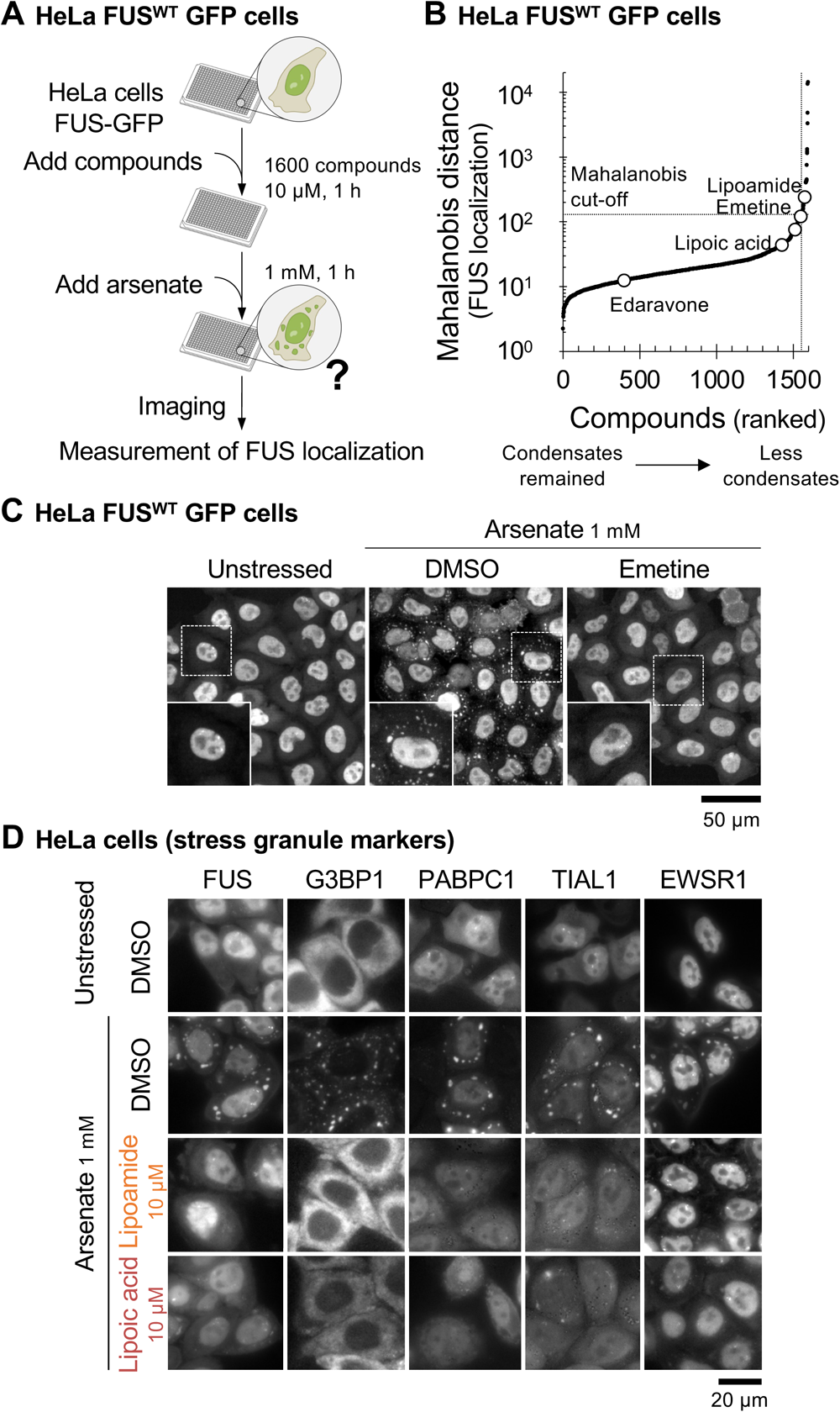
Lipoamide reduces cytoplasmic condensation of stress granule proteins. **A**, Workflow for screening small molecules for effects on FUS-GFP localisàon in HeLa cells *ex vivo*. **B**, Ranked Mahalanobis distances for all 1600 compounds screened (mean from six fields of view) where high values indicate more compound effect. Several automated measures of FUS localisàon were combined into a single Mahalanobis distance score; the largest contributors were cytoplasmic FUS condensate number and area (see the method section). A cut-off of 130 was used to select 47 compounds for further analysis. **C**, The sub-cellular localisàon of FUS-GFP in unstressed HeLa cells, stressed cells with compound solvent (DMSO) negàve control, and with the posìve control emène. Stress causes nuclear export of FUS and formàon of stress granules (cytoplasmic liquid FUS-containing condensates). Insets, magnified images in the boxed areas. **D**, Representàve images of HeLa cells expressing GFP-tagged stress granule markers (G3BP1, PABPC1, TIAL1, or EWSR1) from 3 independent experiments. The cells were pre-treated with 10 μM lipoamide or lipoic acid (with DMSO solvent control) for 1 h followed by 1 mM arsenate for 1 h, or DMSO without arsenate.

The 47 strongest hits in HeLa cells were further tested *in vitro* for an effect on condensàon of purified FUS-GFP under physiological (low salt, 50 mM KCl and reducing, 1 mM DTT) condìons, with the aim of selecting for compounds which can directly affect stress granule proteins (Fig. S1A). Seven compounds significantly affected FUS-GFP condensates *in vitro* and fell into three compound classes (Fig. S1B,C). Of these, surfactants have no plausibility as a systemic therapeùc, and heterotri-and tetracyclic compounds have previously been investigated for anti-prion properties with limited success^25^. Lipoamide was a novel hit for stress granule modulàon. Lipoic acid, a related compound featuring a carboxylic acid instead of the carboxamide, was also a good hit in HeLa cells (Fig. 1B).

### Lipoamide prevents and reverses stress granule formation in cultured cells

To test whether lipoamide and lipoic acid affect stress granule formàon or solely partitioning of FUS into stress granules, we treated HeLa cells expressing five different GFP-tagged stress granule proteins with lipoamide or lipoic acid. Pre-treatment with either compound for 1 h prior to 1 h arsenate stress prevented cytoplasmic condensàon for all proteins we tested, including G3BP1 (Fig. 1D). Addìon of lipoamide and lipoic acid to arsenate-stressed cells, in continued presence of arsenate, also led to dissolùon of pre-existing stress granules (Fig. S1D).

To assess whether lipoamide acts specifically on stress granules, we tested its effects on other intracellular condensates such as P-bodies, Cajal bodies, and DNA damage foci and found that these nuclear or cytoplasmic condensates were not affected (Fig. S1E). The specificity extended to stressor types, as stress granule formàon induced by oxidàve stress and osmòc shock was inhibited whereas stress granules still formed after heat treatment or inhibìon of glycolysis (Fig. S1F). Therefore, we conclude that the lipoamide activity is comparàvely specific with regard to modulàng the properties of a cellular condensate.

To confirm that lipoamide enters cells and to determine its intracellular concentràon, we synthesised [15N]-lipoamide, which can be quantitàvely detected by ^15^N NMR (Fig. S2). Upon treatment of HeLa cells with 100 µM [15N]-lipoamide, loss of [15N]-lipoamide signal from the growth medium indicated that it accumulates in millimolar concentràons in cells. There was also a corresponding gain in [15N]-lipoamide signal in the cell pellet (Fig. S2A–C). These indicate that lipoamide is taken into cells.

We used two strategies to ask whether lipoamide partitions into stress granules. Firstly, we used [15N]-lipoamide with FUS condensates, as a minimal *in vitro* model of the stress granule environment. [15N]-lipoamide from the dilute phase following FUS-GFP condensàon under low salt reducing condìons (Fig. S3A–D) partitioned into the FUS-GFP condensate phase by a factor of ten (Fig. 2A). To analyse partitioning of lipoamide into stress granules in cells, we synthesized a lipoamide analogue derivàsed with a diazirine (for UV-induced crosslinking) and alkyne (for click chemistry) groups (Fig. 2B). This dissolved stress granules with a slightly lower potency than lipoamide (Fig S3E), but allowed us to crosslink this analogue to proteins in the vicinity by UV irradiàon, subsequently labelling it *via* click reaction with a fluorophore. Using the click-crosslink analogue at 30 µM (insufficient to dissolve stress granules), we observed signal particularly in nuclei, mitochondria, and stress granules both without and with crosslinking. Crosslinking increased colocalizàon with stress granules (Figs. 2C,D and S3F). Comparison of signal intensity without and with crosslinking suggests that >50% of the analogues are part of high affinity complexes with fixable macromolecules and/or are covalently bound to proteins in each compartment (Fig. 2E). We suspect that this represents nonspecific binding of the reduced dithiolane to proteins. Taken together, these data indicate that lipoamide partitions into stress granules, along with other organelles.

**Fig. 2.**
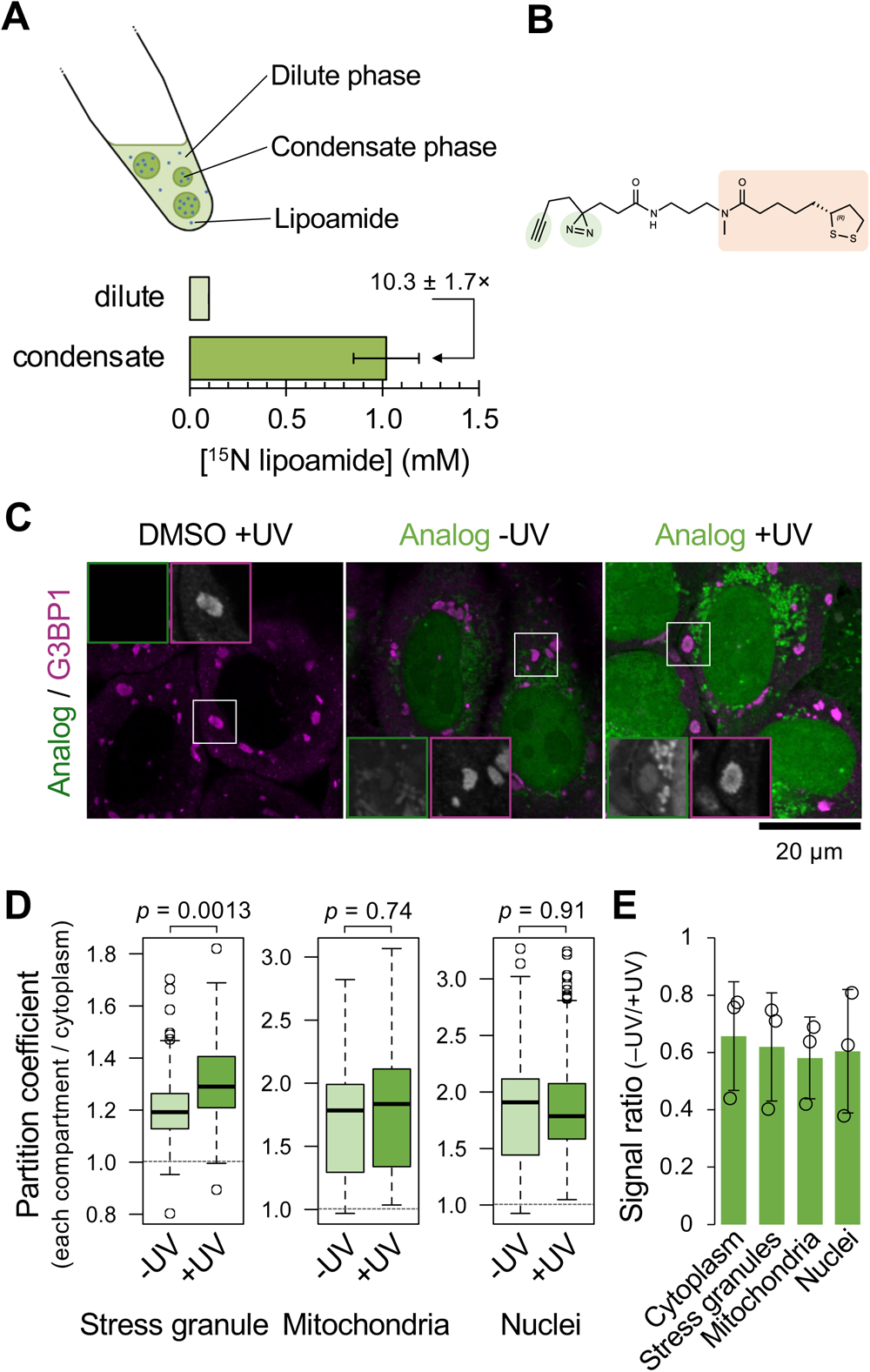
Lipoamide partitions into compartments of stress granule proteins. **A**, Top, schema of lipoamide partitioning into FUS condensates *in vitro*. Bottom, mean ± s.e.m. of concentration of racemic [15N]-lipoamide in the condensate and the surrounding dilute phase of FUS-GFP *in vitro*, quantified using _15_N(^1^H) NMR from 4 independent experiments. **B**, Chemical structure of the click-crosslink lipoamide analog, with the lipoamide backbone (orange) and the groups for UV cross linking and click reaction (green). **C**, Representative images of HeLa cells treated with 3 mM arsenate for 1 h followed by 30 µM of the analogue or the control DMSO in the presence of arsenate for additional 30 min before either irradiated with UV for cross-linking (+UV) or not (-UV), fixed, immunostained, and subjected to the click reaction with the fluorophore (for all conditions). Stress granules were labelled with G3BP1. Insets, stress granules in the boxed areas (analogue and G3BP1 boxed in green and magenta, respectively). **D**, Boxplot of the partition coefficient of the analogue into stress granules, mitochondria, or nuclei relative to the cytoplasm (excluding stress granules and mitochondria) based on signal intensity of the fluorophore. Boxplot shows median (bold bar), 25^th^ and 75^th^ percentiles, and outliers (open dots); whiskers extend to the most extreme values. *n* = 344 (-UV) and 345 (+UV) cells from 3 experiments. *p* values by unpaired *t*-test. **E**, Mean ± s.d. of signal intensity ratio (-UV against +UV) of the fluorophore at indicated subcellular compartments.

### SAR study identifies more potent lipoamide analogues and the dithiolane as a key feature

To determine which chemical features of lipoamide are required for activity, we synthesised a panel of lipoamide-like compounds and tested the structure-activity relàonship (SAR). As a reference, we confirmed lipoamide potency: lipoamide pre-treatment, in both HeLa and induced pluripotent stem cells (iPSCs), caused a dose-dependent decrease in stress granule numbers while an increase in partition of FUS-GFP back to the nucleus with a similar dose-dependency (Fig. 3A). Titràon analyses of the series of lipoamide analogues identified 16 compounds with more than approximately five-fold increased potency (EC_50_ < 2.5 µM, summarised in Fig. 3B–J), compared to lipoamide. Specifically, lipoamide derivàves of 6-amino-3-substituted-4-quinazolinones and five-membered aminoheterocyclic amides represented the most potent analogues (Fig. 3I).

**Fig. 3.**
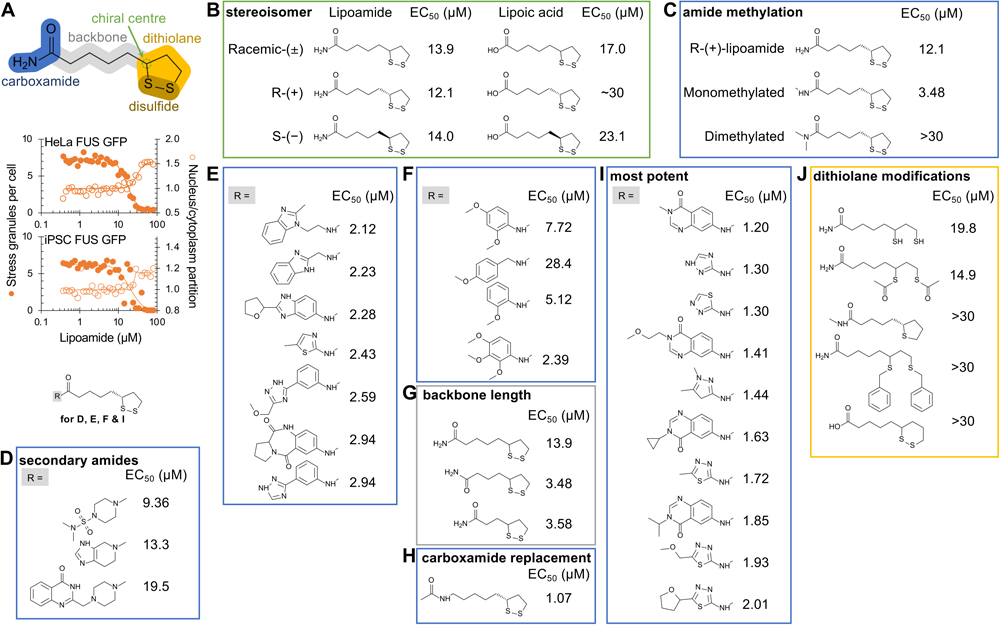
Structure-activity relationship shows lipoamide activity is dependent on the dithiol but is non-enzymatic. **A**, Top, schema of the chemical structure of lipoamide (racemic) with highlighting its features. Bottom, lipoamide dose response using HeLa and iPS cells, showing FUS-GFP condensate (stress granule) number (solid circles, let axis) and nuclear/cytoplasmic signal rào (open circles, right axis) with 1 h pre-treatment with lipoamide followed by 1 h arsenate stress under continued lipoamide treatment. **B-J,** Chemical structures and EC_50_ (µM) of lipoamide and its derivàves, using HeLa cells and the treatment scheme in A. EC_50_ was calculated from dose response curves (details in the method section), and each concentràon of each compound was tested in duplicated wells (*n* = 1750–2650 cells per well) with 2 independent experiments. **B**, Enantiomers of lipoamide and lipoic acid. **C**, Comparison of mono-and di-methylated lipoamide. **D–F**, Addìonal carboxamide analogs of lipoamide. **G**, Modificàons of the linker length between the carboxamide and the dithiolane ring of lipoamide. **H**, Substitùon of the carboxamide of lipoamide. **I**, Carboxamide analogs of 6-amino-3-substituted-4-quinazolinones and five-membered aminoheterocyclic amides. **J**, Modificàons of the dithiolane ring of lipoamide, lipoic acid or similar compounds.

The (*R*) and (*S*) isomers of lipoamide and lipoic acid had a similar EC_50_, indicàng little stereoisomer specificity (Fig. 3B). The chemical structure of lipoamide is similar to that of the lipoyl moiety, used as a hydrogen-accepting cofactor by two mitochondrial Krebs cycle enzymes, which is recycled to the oxidised state by dihydrolipoamide dehydrogenase (DLD, also mitochondrial)^26^. Cells exclusively use the (*R*)-lipoyl moiety stereoisomer. However, the comparable EC_50_ between the (*R*)-and (*S*)-isomers of lipoamide and the absence of a lipoate ligase in eukaryòc cells^26^ indicate that lipoamide does not primarily function through these mitochondrial proteins to effect stress granule dissolùon.

Mono-methylàon of lipoamide on the amide improved the activity, while di-methylàon reduced it (Fig. 3C). However, other disubstituted amide analogues showed activity in the context of more complex amide structures (Fig 3D). Indeed, many mono-substitùons of the lipoamide amide improved the activity, with no clear trend for beneficial substitùons, *i.e.*, a relàvely ’flat’ SAR space in the carboxamide group. Dissimilar substitùons could similarly increase potency to low µM (Fig. 3E), while some similar heterocycle substitùons could have a wide range of potencies (Fig. 3F). Activity could be retained and even increased by shortening the alkane ‘backbone’ (Fig. 3G). A compound without a carboxamide or carboxylic acid moiety (*i.e.*, unlike both lipoamide and lipoic acid) was active, with increased potency (Fig. 3H), although the most potent compounds were mono-substituted amides (Fig. 3I).

Importantly, the dithiolane ring is necessary for activity, indicàng a redox activity for stress granule dissolùon. Lipoamide derivàves are likely reduced in the cellular environment. Indeed, the reduced dihydrolipoamide form was active (Fig. 3J). Furthermore, a labile thiol modificàon (two thioesters) was active while non-labile derivàves (thiol benzylàon and substitùon to a tetrahydrothiophene, a thiolane ring) were not (Fig. 3J). A six-membered disulfide ring removed activity (Fig. 3J). As redox potential is linked to disulfide ring size, this also indicated a redox-linked mechanism. Since Edaravone and ascorbic acid, other redox active compounds^27,28^, did not reduce stress granules at micromolar concentràons comparable to lipoamide (Fig. 1B and S1G), this suggests that lipoamide is more potent than those compounds to control stress granule dynamics. Taken together, the SAR suggests that lipoamide acts through a non-enzymàc route and likely through a redox-associated process.

### Lipoamide weakly increases liquidity of FUS condensates *in vitro*

To test if lipoamide interacts with known stress granule proteins, we turned again to FUS, using the classical methods of isothermal titràon calorimetry (ITC) and chemical shift perturbàon in “fingerprint” ^1^H-^15^N 2D protein NMR spectra. We could not detect interaction of FUS-GFP with lipoamide *in vitro* by ITC. NMR of the N terminal prion-like domain of FUS showed only extremely weak _1_H and ^15^N shifts in the presence of lipoamide (Fig S4A,B). To test if lipoamide alters FUS condensate formàon, we examined the crìcal salt concentràon and temperature of *in vitro* FUS condensates, but found no detectable change in the presence of lipoamide (Fig. S4C).

We then tested whether lipoamide alters FUS condensate properties, first testing the effect on *in vitro* condensate liquidity using laser optical tweezers to assay droplet fusion. This showed significantly decreased droplet fusion times in the presence of lipoamide and thus increased liquidity (higher rào of surface tension to viscosity) (Fig. S4D,E). Over time FUS condensates gradually harden, visible as an increasing viscosity and decreasing mobile fraction of FUS, and eventually forming solid fibres. This is particularly prominent for condensates of ALS-linked mutant FUS G156E, which hardens then forms fibres rapidly^12^. We tested whether lipoamide maintains condensate liquidity, using fluorescence recovery after photobleaching (FRAP). Both lipoamide and lipoic acid reduced FUS G156E-GFP condensate hardening and fibre formàon (Fig. S4F–H).

Finally, we turned to mass spectrometry to analyse changes in FUS G156E self-interaction *in vitro* in the presence of lipoamide. We used lysine-lysine (K-K) chemical crosslinking and, following tryptic digest, mass spectrometry detection of the cross-linked peptides as evidence for inter-and intra-molecular interactions under different condìons. This technique requires lysine residues, which the N-terminal IDR of WT FUS lacks. Therefore, we also analysed a FUS mutant with 12 lysine substitùons in the N-terminal domain. Lipoamide caused a change, predominantly decrease, in the intensity of identified K-K cross-linking sites and therefore suggested reduced FUS-FUS interactions (Fig. S4I,J). Taken together, lipoamide has a weak effect on FUS condensate properties *in vitro* by modulàng FUS-FUS interactions and does so without strong small molecule-protein binding typically detectable by ITC or NMR.

### Lipoamide stabilises proteins with arginine/tyrosine-rich low complexity domains in cells

As the effects of lipoamide on FUS *in vitro* were likely too small to explain the effect of lipoamide on stress granule dynamics in cells, we turned to thermal proteome profiling (TPP). Here, aliquots of HeLa cells treated with DMSO (solvent control), lipoamide, arsenate, or lipoamide and arsenate were heated to a range of different temperatures, and the abundance of soluble proteins was measured by quantitàve mass spectrometry. Relàve increase in abundance with temperature is indicàve of protein thermal stability^29,30^ (Fig. 4A), summarised as z-scores (Fig. 4B and S5A,B). Increased protein thermal stability in the presence of a small molecule often indicates interaction^31,32^. Thermal stabilìes of proteins in lipoamide vs. lipoamide and arsenate-treated cells showed a strong posìve correlàon, but arsenate vs. lipoamide and arsenate-treated cells did not, indicàng a dominant effect of lipoamide. Furthermore, lipoamide treatment also broadly reversed the thermal stability changes occurring due to arsenate treatment (Fig. S5A,B). Therefore, we focused on the analysis of the sample treated with both lipoamide and arsenate. As a posìve control, we confirmed that the thermal stability of DLD was weakly but significantly increased, consistent with lipoamide binding to the active site (z = 0.66±0.007, adjusted *p*-value: false discovery rate [FDR] = 2.6 × 10^−4^). As we would predict from its mitochondrial localizàon and enzymàc function, RNAi of DLD affected neither stress granule formàon nor the lipoamide activity to prevent it (Fig. S5C). Histone deacetylase 1 (HDAC1) and HDAC2 were also stabilized (z = 3.58, FDR = 1.8 × 10^−14^ and z = 4.75, FDR = 7.4 × 10^−5^, respectively), consistent with a recent report^33^.

**Fig. 4.**
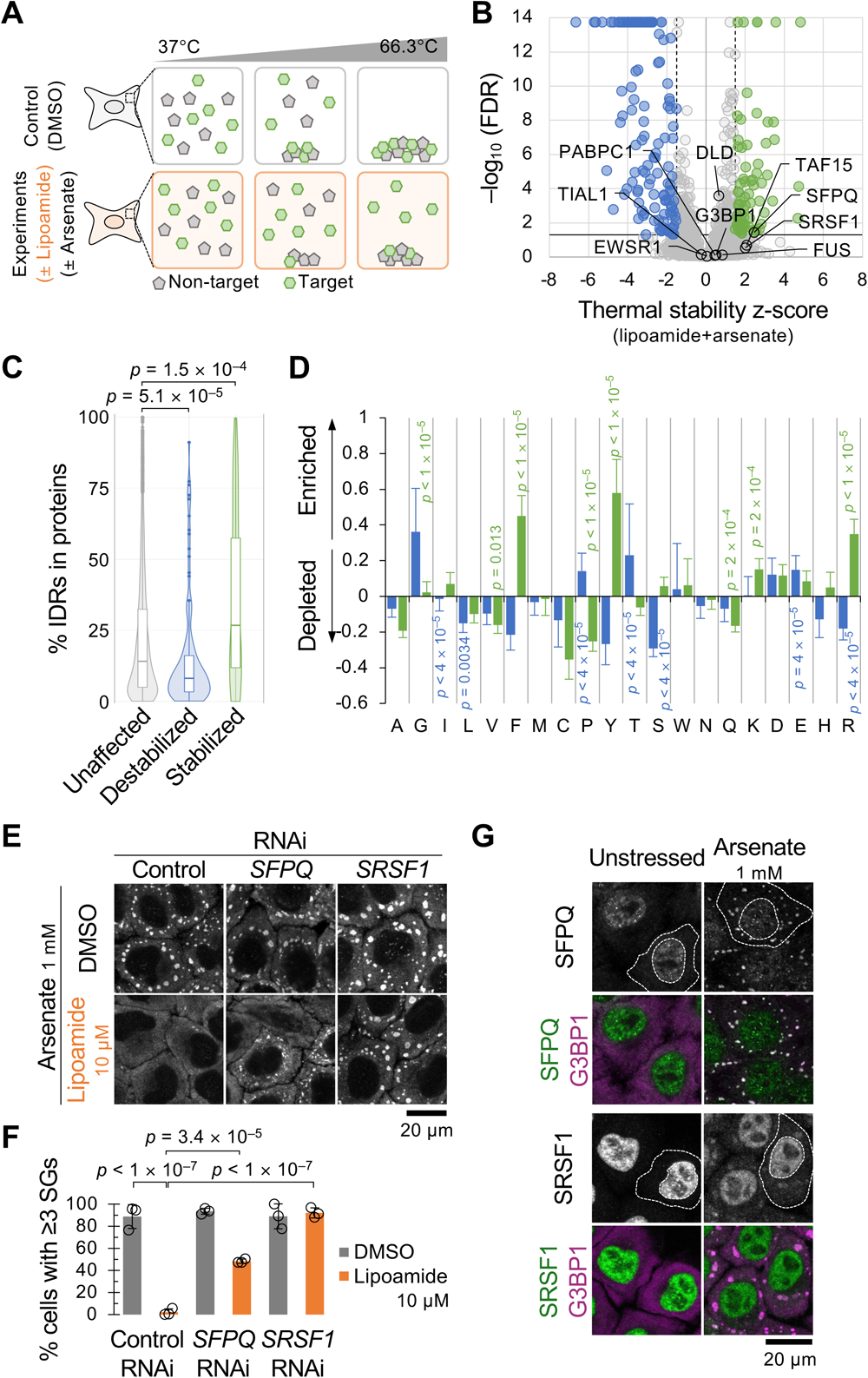
Lipoamide interacts with disordered proteins in cells. **A**, Schema of TPP to see the effect of lipoamide treatment on protein thermal stability. In our TPP, HeLa cells treated with 100 µM lipoamide and/or 1 mM arsenate were heated to ten different temperatures. Heàng causes protein denaturing and precipitàon, and lipoamide could prevent precipitàon of the target proteins. The soluble protein amount was quantified by mass spectrometry and normalized with the result of non-stressed 0.1% DMSO-treated samples (control). **B**, Volcano plot of z-scores (mean from 3 individual experiments) and FDRs of protein thermal stability in HeLa cells treated with lipoamide and arsenate. A larger z-score indicates more thermal stabilizàon. Black broken and solid lines indicate cutoffs of z-score (±1.5) and FDR (<0.05), respectively, used to classify stabilized (green) and destabilized (blue) proteins in F and G. The posìons of DLD, SFPQ, SRSF1, and several stress granule proteins are indicated. **C**, Violin and box plots showing proportions of IDRs in each protein from cells treated with both lipoamide and arsenate. The proteins were categorized into stabilised (z > 1.5, FDR < 0.05; 70 proteins), destabilised (z < –1.5, FDR < 0.05; 144 proteins), and unaffected (the others detected; 5811 proteins). Boxplots show median (bold bar), 25^th^ and 75^th^ percentiles, and outliers (closed dots); whiskers extend to the most extreme values. *p* values by a Wilcoxon signed-rank test followed by Holm’s test. **D**, Enrichment (> 0) or deplèon (< 0) of each amino acid in IDRs of the stabilized (green) and destabilized (blue) proteins in cells treated with both lipoamide and arsenate, in comparison to IDRs of all the detected proteins as background. *p* values by unpaired *t*-test followed by Bonferroni’s test. **E**, Representàve images of HeLa cells from >3 independent experiments, depleted of SFPQ or SRSF1, treated with 10 µM lipoamide or 0.1% DMSO for 1 h followed by 1 mM arsenate for 1 h in the presence of lipoamide. Stress granules were labelled with G3BP1. **F**, Mean ± s.d. of percentage of stressed HeLa cells with ≥3 G3BP1-posìve stress granules (SGs). *n* = 292–615 cells from 3 independent experiments. Dots indicate means of each experiment. *p* values by Tukey’s test. **G**, Representàve images of HeLa cells from >3 independent experiments, treated with 1 mM arsenate for 1 h and subjected to immunostaning with indicated antibodies. Outer and inner broken lines indicate edges of the cytoplasm and nucleus of one cell each condìon, respectively. Note that the SFPQ and SRSF1 signals were diminished by individual RNAis.

Many proteins had higher TPP z-scores than DLD (Fig. 4B): Lipoamide and arsenate treatment resulted in significantly increased thermal stability of 70 proteins, while reducing the thermal stability of 144 proteins compared to no treatment (Fig. 4B). Stabilised proteins had disproportionately long IDRs which contained a disproportionally high proportion of arginine (R), tyrosine (Y), and phenylalanine (F) residues, while destabilised proteins showed the opposite trend (Fig. 4B–D). R and Y-rich IDRs are characteristic of stress granule proteins such as the FET family (FUS, TAF15, and EWSR1)^34^, although their thermal stability was not significantly increased (z = 0.84±1.3, FDR = 0.73; z = 2.08±1.5, FDR = 0.20; z = 0.05±0.82, FDR = 0.87, respectively). This is consistent with no obvious interaction between FUS and lipoamide *in vitro*. Also, individual FET family proteins are neither necessary for stress granule formàon^10^ nor lipoamide activity (Fig. S5C), further indicàng that they are not primary targets of lipoamide for stress granule dissolùon.

### Specific stress granule proteins are necessary for lipoamide activity in cells

To assess which of the proteins identified as interacting with lipoamide by TPP are necessary for lipoamide activity, we performed an endoribonuclease-prepared small interfering RNA (esiRNA)^35^-mediated gene knockdown screen of all 122 proteins with increased thermal stability (z > 2) (Table S1, Fig S5A). We looked for the genes whose deplèon reduced lipoamide activity in preventing stress granule formàon, which identified two IDR-rich proteins: splicing factor proline-and glutamine-rich (SFPQ) and splicing factor serine/arginine-rich splicing factor 1 (SRSF1)^36,37^: Lipoamide pre-treatment failed to prevent stress granule formàon in cells with either gene deplèon (Figs. 4E,F and S5D). RNAi of SFPQ or SRSF1 also prevented dissolùon of pre-existing stress granules following lipoamide treatment (Fig. S5E). Stress granule formàon was neither exacerbated in stressed cells nor induced in non-stressed cells by these RNAis, showing that the phenotype of lipoamide pre-treated cells is not simply due to basal increase in stress granule formàon (Figs. 4E,F and S5F).

Both SFPQ and SRSF1 are stress granule proteins: like FUS and TDP-43, both localise to the nucleus in unstressed cells and, in stressed cells, stress granules (Fig. 4G). Therefore, lipoamide activity dissolving stress granules is dependent on at least two stress granule proteins.

### Redox state controls SPFQ-mediated dissolution of stress granule protein condensates

The activity of lipoamide requires the redox active dithiolane (Fig. 3J), and SFPQ is notably rich in redox-sensìve methionine (28 out of 707 amino acids; Fig. 5A). SRSF1 is not methionine-rich with only three out of 248 amino acids. Pioneering work has previously shown that methionine oxidàon modulates function and material properties of phase separated yeast ataxin-2^38^, and methionine oxidàon in SFPQ has been detected in cells^39^. These suggest that SFPQ may be a main target of lipoamide in a redox-based mechanism of action. We used in vitro experiments using pure protein to analyse the effect of oxidàon of SFPQ on condensate formàon, using the oxidizing agent hydrogen peroxide (H_2_O_2_). SFPQ condensàon was induced at low salt concentràon (75 mM KCl; Fig. S6A). Oxidàon of SFPQ, confirmed by modulated electrophorèc migratory aptitude in non-reducing SDS-PAGE (Fig. S6B), led to dissolùon of SFPQ condensates in an H_2_O_2_ concentràon-dependent manner (Fig. S6A), similar to the behavior of axaxin-2 condensates^38^. This suggests that oxidàon alters properties of SFPQ proteins. In contrast, H_2_O_2_ alone did not lead to FUS condensate dissolùon (Fig. S6A). Therefore, oxidàon-mediated condensate dissolùon is specific to a subset of proteins.

**Fig. 5.**
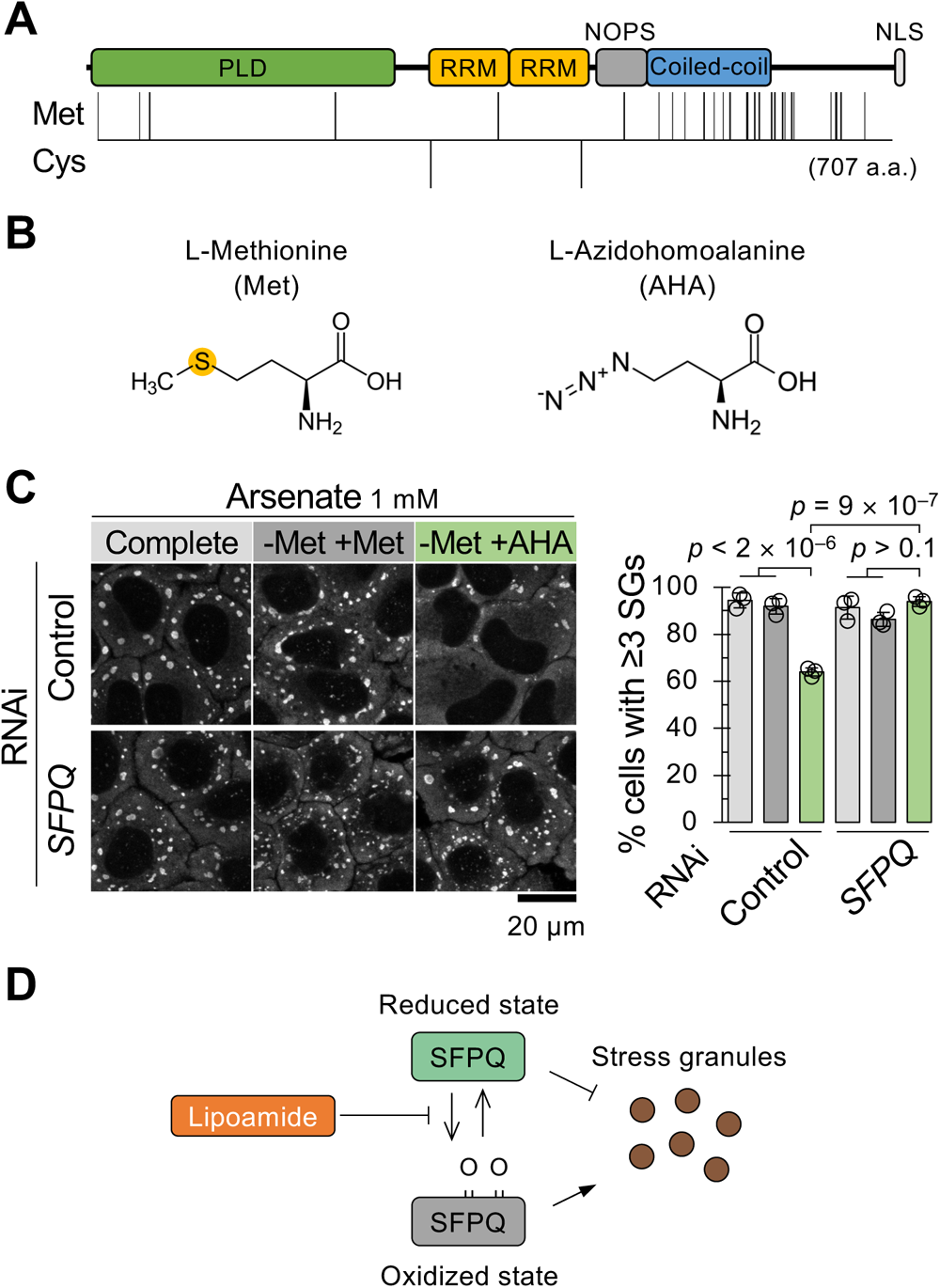
SFPQ redox state may mediate the lipoamide activity. **A**, Schema of distributions of methionine (Met; 28 residues) and cysteine (Cys; 2) residues in human SFPQ. PLD, prion-like domain; RRM, RNA recognition motif; NOPS, NonA/paraspeckle domain; NLS, nuclear localizing signal. **B**, Chemical structures of Met and its non-natural analogue azidohomoalanine (AHA). **C**, Left, representative images of HeLa cells subjected to indicated RNAis, cultured in complete medium (light grey) or Met-free medium supplemented with 1 mM of Met (dark grey) or AHA (green) for 2 h followed by 1 mM arsenate for 1 h (experimental schema in Fig. S6E). Stress granules (SGs) were labelled with G3BP1. Right, mean ± s.d. of percentage of stressed HeLa cells with ≥3 G3BP1-positive SGs. *n* = 325–407 cells from 3 independent experiments. *p* values by Tukey’s test. **D**, Schema of SFPQ as a redox sensor to modulate stress granule condensation.

We next examined functions of SFPQ and its oxidàon on FUS condensate dynamics at a physiological salt concentràon (150 mM KCl), which kept SFPQ proteins in diffused state (Fig. S6C). We found that adding SFPQ prevented FUS condensàon (Fig. S6C). This effect did not occur by adding only GFP proteins (Fig. S6C). However, H_2_O_2_ treatment restored FUS condensates even in the presence of SFPQ (Fig. S6D). These suggest that SFPQ proteins dissolve stress granule protein condensates in a redox state-dependent manner.

Based on these *in vitro* results, we hypothesized that stress granule formàon would be attenuated if SPFQ is not oxidizable. To assess this possibility, we aimed to replace methionine with a non-oxidizable, non-natural analogue L-azidohomoalanine (AHA), normally used for protein labeling^40^ (Fig. 5B). Cells were cultured in methionine-free medium supplemented with AHA for 2 h, resulting in methionine-to-AHA replacement in newly synthesized proteins, and then stressed with arsenate for 1 h (Fig. S6E). This resulted in attenuated stress granule formàon (Fig. 5C), and normal stress granule formàon was rescued by deplèon of SFPQ (Fig. 5C). This suggests that SFPQ in the reduced state is responsible preventing stress granule formàon. One possibility is that cellular stress leads to oxidàon of SFPQ, allowing stress granule condensàon. In this scenario, lipoamide reduces SFPQ and restores its stress granule dissolùon activity (Fig. 5D).

### Lipoamide treatment rescues nuclear localization and functions of FUS and TDP-43

FUS and TDP-43 have important nuclear functions in unstressed cells. We asked whether lipoamide treatment not only dissolves stress granules but also returns these proteins to the nucleus. Similar to FUS-GFP (Fig. 3A), lipoamide pre-treatment increased partition of TDP-43 and the ALS-associated nuclear localisàon sequence mutant FUS P525L-GFP to the nucleus in stressed HeLa cells (Fig. S7A). To confirm that these effects also occur in cells prominently defective in ALS, we analysed iPSC-derived motor neurons (MNs). Lipoamide had similar effect on nuclear partitioning of wild-type TDP-43 in stressed (prolonged oxidàve stress with low dose [10 µM] of arsenite) and FUS P525L-GFP in non-stressed but long-cultured condìons (Figs. 6A,B and S7B,C)^41^.

**Fig. 6.**
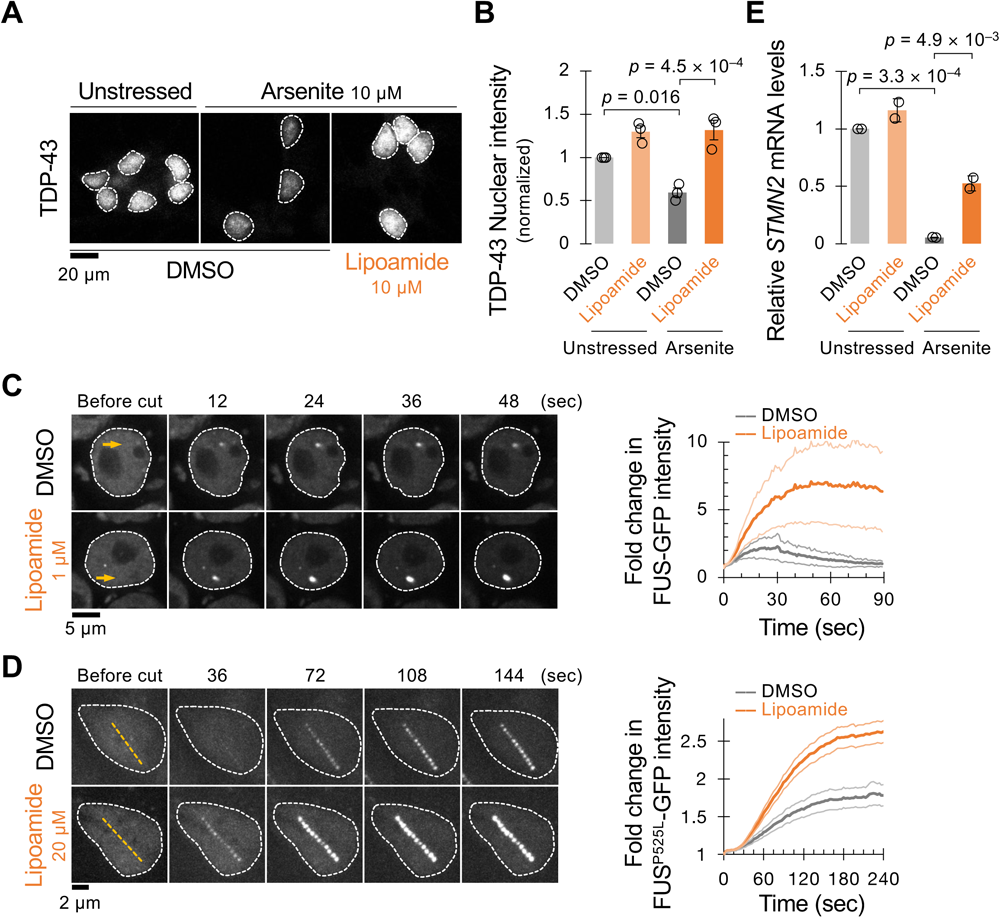
Lipoamide improves nuclear localization of FUS and TDP-43. **A**, Representàve images of iPSC-derived MNs from 3 independent experiments, treated with 0.1% DMSO or 10 µM lipoamide for 1 day followed by 10 µM arsenite for 5 days in the presence or absence of lipoamide, and labelled with TDP-43. Broken line, outline of nuclei. **B**, mean ± s.e.m of nuclear TDP-43 levels normalized to those of unstressed DMSO-treated MNs (control). *n* = 417–1741 cells from 3 independent experiments. *p*-values, Tukey’s test. **C**, (Let) images showing recruitment of FUS-GFP to sites of UV laser-induced DNA damage (yellow arrow) in nuclei (outlined with broken lines) of iPS cells at indicated times after laser irradiàon. Cells were subjected after 1 h treatment with lipoamide followed by 1 h arsenate stress. (Right) mean ± s.d. of relàve FUS-GFP signal intensity in response to DNA damage. *n* = 5 (DMSO) and 7 (lipoamide) cells. **D**, (Let) images of nuclei (outlined with broken lines) of iPSC-derived MNs expressing FUS P525L-GFP from 3 independent experiments, cultured for 21 days and then treated with 0.02 % DMSO or 20 µM lipoamide for 24 h, at indicated times after laser irradiàon. Yellow lines indicate laser-irradiated sites. (Right) mean ± s.e.m. of relàve intensity of FUS-GFP at DNA damage sites after ablàon. *n* = 14 (DMSO) and 18 (lipoamide) cells from 3 independent experiments. **E**, Mean ± s.d. of relàve STMN2 full length mRNA levels normalized to those of GAPDH from 2 independent experiments. *P*-values by Tukey’s test. In B and E, MNs were treated as in A.

We characterised the functional importance of the re-localizàon to the nucleus by considering FUS and TDP-43 nuclear functions. FUS forms condensates at DNA damage sites to engage in DNA damage repair, and this malfunction caused by ALS-linked mutàons on FUS is implicated to underlie neuronal dysfunction in ALS^42^. Lipoamide increased recruitment of FUS-GFP (WT in iPSCs and P525L in iPSC-derived MNs) to laser-induced DNA damage foci (Fig. 6C,D). TDP-43 contributes to normal transcript splicing in the nucleus, particularly of Stathmin-2 (STMN2) transcript, and altered *STMN2* splicing leading to reduced transcript levels is a hallmark of ALS^6,43^. In iPSC-derived MNs, the prolonged oxidàve stress recapitulated reduction in *STMN2* mRNA levels. This reduction in splicing was rescued by lipoamide treatment, concomitant with TDP-43 nuclear partitioning (Fig. 6B,E). Lipoamide action therefore dissolves stress granules, allows return of those ALS-linked proteins to the nucleus, and restores nuclear functions of FUS and TDP-43.

### Lipoamide alleviates ALS phenotypes in familial ALS models

The ultimately lethal phenotype of ALS is thought to be caused by axon defects in motor neurons. Indeed, iPSC-derived MNs expressing FUS P525L show a motor neuron survival defect *in vitro*, with reduced neurite growth and defective axonal transport^41^. Lipoamide treatment rescued neurite growth of iPSC-derived MNs stressed with arsenite, shown by increased area covered in neurites in a non-polarised culture (Fig. 7A). We tested if this correlated with improved axonal transport, by tracking lysosome transport in iPSC-derived MN axons grown through silicone channels (Fig. 7B). As previously observed, in an unstressed condìon, distal axonal transport of lysosomes was disrupted by expression of FUS P525L^41^, and lipoamide recovered transport to a similar level to that in WT FUS iPSC-derived MNs (Fig. 7C,D). Motor neuron degeneràon caused by an ALS-associated FUS mutant can therefore be rescued by lipoamide.

**Fig. 7.**
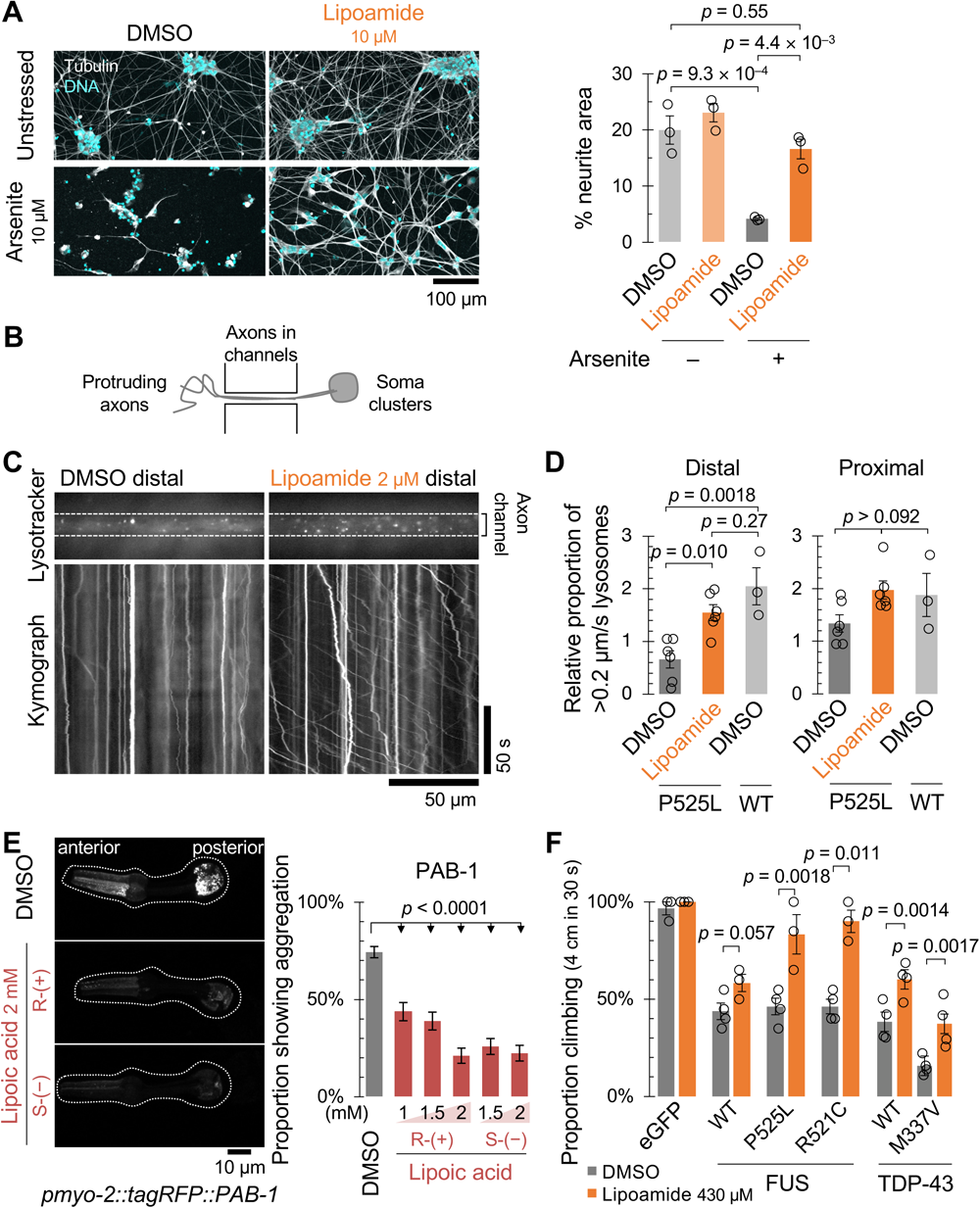
Lipoamide improves cellular fitness in ALS disease models of iPSC-derived motor neurons and animals. **A**, (Let) representàve images of iPSC-derived MNs treated as in Fig.5A. (Right) mean ± s.e.m of percentage of neurite (tubulin-posìve) area. 18 image fields from 3 independent experiments. *P*-values, Tukey’s test. **B**, Schemàc of neuron culture, showing the channels through which the axons grow from the soma on the right. **C**, Kymographs of lysosome movement in the distal portion of FUS P525L MN axons 3 days after treatment with compound solvent (DMSO) or 2 μM lipoamide, visualized with lysotracker. **D**, mean ± s.e.m. of relàve proportion of lysotracker-labelled lysosomes moving with an average speed greater than 0.2 μm/s following 3 days treatment with 2 μM lipoamide or equivalent DMSO concentràon solvent control for iPSC-derived MNs expressing either P525L or WT FUS, normalized to mean of proportion moving (proximal and distal) in the DMSO-treated P525L FUS MNs. 6 (P525L) or 3 (wild-type) biological replicates, analyzing 5 axon bundles per replicate. *p* values by Tukey’s test. **E**, (Let) Representàve images of the pharynx of worms expressing fluorescently tagged PAB-1 with or without lipoic acid treatment (2 mM). Broken lines, the edge of pharynges. (Right) Mean ± s.e.m. of incidence of each protein aggregàon in the pharyngeal muscles. Incidence of PAB-1 aggregàon was scored from the proportion of animals with >10 aggregates. \*\*\**p* < 0.0001 by Fisher’s exact test. *n* > 100 for each sample. **F**, Mean ± s.e.m. of proportion of flies that climbed, fed with 0.1% DMSO (solvent control) or 430 µM lipoamide. Human WT or ALS-linked mutants of FUS (let) or TDP-43 (right) were expressed in motor neurons. *p* values by unpaired *t*-test. *n* = 30–40 (FUS) and 130–202 (TDP-43) flies from 3 or 4 independent experiments.

Aggregàon of TDP-43 and FUS in neurons is a hallmark pathology of ALS, and aggregàon of proteins is also a phenotype of aging more generally, including in *C. elegans*^44^. Feeding lipoic acid (it has higher solubility in food media than lipoamide) caused a dose-dependent reduction in the number of aggregates of transgenic PAB-1, an orthologue of the human stress granule protein PABPC1 (Fig. 7E), but not those of a non-stress granule protein RHO-1 (Fig. S8A)

In *D. melanogaster*, motor neuron-specific expression of human FUS and TDP-43 induces ALS-like phenotypes, including motor defects manifesting as a reduced ability for negàve geotaxis^45,46^. Feeding either lipoamide or lipoic acid improved climbing ability in flies expressing FUS nuclear localizing signal (NLS) mutants, either FUS P525L or R521C (Figs. 7F and S8B). Similarly, lipoamide feeding alleviated climbing defects in flies expressing TDP-43, either WT or an ALS-linked mutant M337V (Fig. 7F). The severe phenotype caused by TDP-43 M337V was associated with abnormal neuromuscular junction morphology, the presence of satellite boutons, similar to previously described phenotypes of another ALS-linked mutant TDP-43 G298S^47^. Lipoamide treatment suppressed appearance of satellite boutons (Fig. S8C). Collectively, our data show that lipoamide can alleviate ALS-like phenotypes in pàent-derived motor neurons and animal models caused by expression of two ALS-associated stress granule protein mutants.

## Discussion

Stress granules are an example of a liquid cellular compartment formed by phase separàon. Due to the strong genèc associàon of ALS with stress granule proteins, we sought small molecules which alter the physiological function of stress granule proteins in forming biological condensates. Our screen identified lipoamide, which partitioned into stress granules in cells and their *in vitro* model (condensates formed with the stress granule protein FUS). Lipoamide caused rapid disassembly and prevented formàon of stress granules. Our SAR showed the potency of lipoamide for stress granule dissolùon could be increased, particularly by mono-substituted amines and some alkane backbone modificàons.

The dithiolane is always required for activity of lipoamide-like molecules. However, the degree to which other areas of lipoamide could be modified while retaining activity was large. This shows the lipoamide compound family has potential for medicinal chemistry development. The high degree to which the non-dithiolane regions could be altered would be surprising for a molecule that binds a protein at a structured binding site, although this is perhaps unsurprising given the interaction of lipoamide with intrinsically disordered proteins. Overall, lipoamide is likely therefore acting by delivering a redox-active dithiolane payload to physicochemical environments formed by its interacting proteins, including stress granules. More active lipoamide derivàves are likely improving targèng (cell uptake and partition into stress granules) while leaving the dithiolane payload intact.

Our work has identified a plausible candidate for the protein likely to sense the redox state of the cell. Among all the proteins stabilised by lipoamide, our RNAi screen showed only SFPQ and SRSF1 were necessary for the rapid (<20 minute) lipoamide effect on stress granule dynamics. Both are stress granule proteins. Uniquely, SFPQ is very methionine-rich, which likely confers high redox sensìvity not found in most arginine/tyrosine-rich IDR-containing proteins. SFPQ activity was indeed methionine-and redox-dependent. SFPQ appears the primary target for early lipoamide activity, which overrides the ability of SFPQ to act as an oxidàon sensor, promòng stress granule dissolùon only when reduced. While we saw some effect of lipoamide on FUS condensate liquidity *in vitro*, we suspect that this is a similar effect of secondary importance on a less redox-sensìve stress granule protein or a minor effect from strong partitioning of lipoamide into this condensate. Overall, our identificàon of lipoamide allowed us to dissect a mechanism where dissolùon of stress granule condensates involves direct redox sensìvity of a key stress granule protein. Lipoamide therefore represents productive small molecule intervention in an emerging paradigm: redox-sensìvity of proteins able to phase separate as a homeostàc mechanism^38,48,49^.

Oxidàve stress is a common theme in ALS pathogenesis mechanisms^50–52^. We showed that lipoamide can recover pathology in motor neurons and animals expressing ALS-associated stress granule mutants (FUS and TDP43) with no explicit oxidàve stress. This leaves the link between the redox-associated cellular effects of lipoamide on stress granules and neuron/animal model outcomes ambiguous, but consistent with modulated stress granule formàon in response to stochastic oxidàve stresses. Previous works described that HDAC1 and 2 are responsible for long-term changes in histone acetylàon in motor neurons^53^ and that they are inhibited by lipoamide^33^. However, we did not find them necessary for short-term lipoamide activity on stress granules, instead detecting the two stress granule proteins both necessary for lipoamide activity. However, this does not preclude long-term nuclear effects. FUS NLS mutàons are strongly associated with ALS^41,54–56^ and dissolùon of stress granules by lipoamide leads to return of FUS and TDP-43 to the nucleus. Indeed, we saw lipoamide does not prevent nuclear FUS condensàon and rescues nuclear TDP-43 functions. Ultimately, this relates to central questions about ALS pathogenesis. Does persistent stress granules or stress granule protein aggregàon lead to harmful (gain-of-function) effects, or is stress granule formàon fundamentally beneficial over a short time scale but, over a long time scale, leads to defects from nuclear loss-of-function by sequestràon of proteins in the stress granules? – overall our results are consistent with the latter.

Although lipoamide does not have the characteristics of a typical therapeùc, it is notable that lipoic acid is used to treat diabèc neuropathy and, in humans, a 600 mg daily dose gives plasma concentràons of 8 to 30 μM^57,58^. This is comparable to the concentràons used in our cell-based assays and we saw beneficial effects in pàent-derived motor neurons and *D. melanogaster* models of ALS. Therefore lipoamide, in addìon to allowing our discovery of direct redox sensàon by SFPQ for stress granule dissolùon, has some plausibility as the basis of a therapeùc with medicinal chemistry potential for further improvements. However it is important to point out that we have not shown a direct relàonship between stress granule dissolùon and phenotype rescue on our disease models. Future work will be required to understand the relàonship between stress granule formàon and disease in animal models. However the identificàon of lipoamide provides a powerful tool for such investigàon.

## Methods

### Cells and cell lines

Kyoto HeLa cells were maintained in high glucose Dulbecco’s modified Eagle’s medium (DMEM, Thermo Fisher Scientific) supplemented with 10% FBS and 1% penicillin-streptomycin at 37°C with 5% CO_2_. Stable Kyoto HeLa BAC cell lines expressing proteins with a C-terminal fluorescent protein were generated using BAC recombineering^59^. This gives near-endogenous expression levels of the fusion protein^12,60^. In these lines, GFP is part of a modified localisàon and affinity purificàon (LAP) tag^61^, providing a short linker. The stable expression was kept under G418 selection (400 μg/ml; Gibco). The following BAC lines were used: FUS (MCB_005340) (used also for the compound screen [see Small molecule screen]), COIL (MCB_0002582), DCP1A (MCB_0003876), EWSR1 (MCB_0008863), PABPC1 (MCB_0006901), TIAL1 (MCB_0008967), and TRP53BP1 (MCB_0003740). HeLa FUS P525L-GFP cells were generated similarly to the iPS cell lines as described previously^41^.

Human iPS cells were grown in either TeSR E8 or mTeSR1 medium (Stem Cell Technologies) at 37°C with 5% CO_2_ (ref.^62^). iPS cells lines, derived from two different donors, expressing FUS with a C-terminal GFP fluorescent marker were used. All were generated using CRISPR/Cas9 assisted tagging and mutagenesis and have been previously described^41^. KOLF iPS cell lines expressing wild-type FUS-GFP or FUS P525L-GFP were previously generated from the KOLF-C1 clonal iPS cell line produced as part of the human induced pluripotent stem cell inìàve (HipSci)^63^. KOLF-C1 cells were derived from a healthy male donor. In these lines, GFP is part of a modified localisàon and affinity purificàon (LAP) tag^61^, giving an identical fusion protein sequence to the Kyoto HeLa BAC cell line. AH-ALS1-F58 iPS cell expressing FUS P525L with a C-terminal GFP fluorescent marker were previously generated from a clonal iPS cell line from a female ALS pàent expressing FUS P521C. The P525L mutàon and GFP tag were introduced and the P521C mutàon corrected by simultaneous tagging and mutagenesis^41,64,65^.

MNs for FUS P525L dynamics were induced and maintained as described previously^66^. MNs for the prolonged arsenite stress assay were derived from commercially available WTC-11 iPS cells (Coriell Institute GM25256) and differentiated as described previously^67^. MNs for axonal lysosome mobility assays were generated from AH-ALS1-F58 iPS cells expressing FUS P525L. In short, the iPS cells were differentiated into neuronal progenitor cells and matured to spinal MNs in Matrigel-coated plates as previously described^41,62^. The coàng and assembly of silicone microfluidic chambers (MFC; RD900, Xona Microfluidics) to prepare for the subsequent seeding of MNs was performed as described previously^41,68,69^. MNs were eventually seeded into one side of an MFC for maturàon to obtain a fully compartmentalized culture with proximally clustered somata and their dendrites being physically separated from their distal axons, as only the latter type of neurite was able to grow from the proximal seeding site through a microgroove barrier of 900 µm-long microchannels to the distal site (Fig. 7B). All subsequent imaging in MFCs was performed at day 21 of axonal growth and MN maturàon (day 0 = day of seeding into MFCs).

All procedures using human cell samples were in accordance with the Helsinki convention and approved by the Ethical Committee of the Technische Universität Dresden (EK45022009, EK393122012).

### Recombinant protein purification

Recombinant proteins were purified using baculovirus/insect cell expression system, as previously described^12^. Briefly, 6×His-MBP-FUS-GFP and 6×His-MBP-FUS-SNAP were purified from Sf9 cell lysate by Ni-NTA (QIAGEN) affinity purificàon. The 6×His-MBP tag was cleaved by 3C protease, concentrated by dialysis, and further purified by size exclusion chromatography. 6xHis-MBP-SFPQ-GFP was purified from Sf9 cell lysate by affinity purificàon using Ni-NTA and amylose resin (New England Biolabs). The 6×His-MBP tag and, if necessary, the GFP tag were cleaved by 3C protease and TEV protease, respectively, and the target proteins were concentrated by dialysis and further purified by càon exchange chromatography. The composìon of the storage buffer for the purified proteins was 1 M or 500 mM KCl, 50 mM Tris-HCl pH 7.4, 5% glycerol and 1 mM DTT, and FUS concentràon was adjusted to 30 μM in storage buffer prior to use.

### Small molecule screen

For the small molecule screen we used the PHARMAKON 1600 library of small molecules, prepared as 10 mM solùons in DMSO. The Kyoto HeLa BAC cell line stably expressing FUS-GFP was seeded at 4000 cells per well in 384 well plates 24 h before the assay. The cells were pre-treated with 10 μM compound for 1 h and then stressed with 1 mM potassium arsenate (A6631, Sigma Aldrich). After 1 h, cells were fixed in 4% formaldehyde and stained with 1 µg/ml Hoechst 33342 and CellMask blue (1:10,000; H32720, Thermo Fisher Scientific) before imaged on a CellVoyager CV7000 automated spinning disc confocal microscope (Yokogawa) with a 40× NA 1.1 air objective to assess FUS-GFP localisàon.

FUS-GFP signal was analysed using CellProfiler^70^, and the data were processed with KNIME. Cytoplasm and nuclei were distinguished with weak (CellMask blue) and strong (Hoechst 33342) blue fluorescent signals, respectively. Particle number and sum area, granularity (at 9, 10, and 11 px in the cytoplasm or 1, 5, 6, 7, 8, and 9 px in the nucleus) scale, texture (at 10 px scale), and integrated signal intensity of FUS-GFP in the nucleus and cytoplasm were measured. Z scores (*z* = (*x* − *μ*)/*σ* where *x* is the observed value, *μ* the control mean and *σ* the control standard deviàon) relàve to the DMSO treated control wells on each plate were calculated for these parameters and combined into the Mahalanobis distance. Compounds of interest were selected on the criteria of: treatment returned the cells to the unstressed state (ie. reducing stress granule number, increasing nuclear signal), a clear monotonic dose dependent response, and by manual priorìsàon by known mechanism (e.g. emène, cardiac glycosides) or implausibility as a cell-compàble compound (e.g. surfactants, used as topical antiseptics).

The follow-up *in vitro* assay of compounds on FUS-GFP condensates was assessed in a 384 well plate format. The compound volume (in DMSO) necessary for 1, 3, 10, 30 or 100 μM final concentràon were added by acoustic dispensing (Labcyte Echo 550) to 96 well plate wells containing FUS-GFP in 3 µl of 50 mM Tris-HCl pH 7.4, 1 mM DTT, and 170 mM KCl. Final DMSO concentràon was 0.01 to 1%. Using a Freedom Evo 200 liquid handling workstàon (TECAN) the FUS-GFP/compound mixture was diluted in 7 μl 50 mM Tris-HCl pH 7.4 to reach the final composìon of 50 mM Tris-HCl pH 7.4, 1 mM DTT, 50 mM KCl, the indicated concentràons of each compound and DMSO, and 0.7 μM FUS-GFP. Compound/FUS-GFP and assay buffer were mixed by a standardised pipeãng procedure, split to four wells in clear bottom 384 well plates, and then immediately imaged using a CellVoyager CV7000 automated spinning disc confocal microscope (as above). Condensates in suspension for six fields of view were imaged as a maximum intensity projection of 6 focal planes at 2 μm steps per sample. Condensate number and FUS-GFP partition into condensates were analysed with a fixed intensity threshold using Fiji. Number of condensates and partition were weakly time dependent due to condensate sedimentàon, so normalised assuming a linear change over time by reference to DMSO controls at the start and end of each plate row.

### Compound characterisation on cells

Compound effects were assessed under a variety of condìons in HeLa cells, iPS cells, or iPSC-derived motor neurons. Unless otherwise indicated, cells were pre-treated for 1 h using 10 μM compounds from 10 mM stock in DMSO (or an equal volume of DMSO control) then stressed for 1 h with 1 mM potassium arsenate still in the presence of the compounds. Live cells were imaged by widefield epifluorescence using an inverted Olympus IX71 microscope with a 100× NA 1.4 Plan Apo oil immersion objective (Olympus) and a CoolSNAP HQ CCD camera (Photometrics), using a DeltaVision climate control unit (37°C, 5% CO_2_) (Applied Precision).

Various cellular stresses were achieved by replacing 1 h 1 mM potassium arsenate treatment with other condìons: 0.4 M sorbitol (S1876, Sigma Aldrich) from a 4 M stock in water for 1 h (osmòc stress); 42°C in normal growth medium for 30 min (heat stress); 100 mM 6-deoxyglucose (D9761, Sigma Aldrich) from a 1 M stock in H_2_O in glucose free DMEM (11966025, Thermo Fisher Scientific) supplemented with 10% FCS for 1 h (glycolysis stress). L-ascorbic acid (A4544, Sigma Aldrich) was used from 1 M stock in H_2_O. Sodium arsenite (S7400, Sigma Aldrich) was used from 10 mM stock in H_2_O.

### Compound dose responses

Dose dependent effect of lipoamide on HeLa and iPS cells expressing FUS-GFP were assessed with pre-treatment of lipoamide for 1 h followed by 1 h treatment with 1 mM potassium arsenate similar to the *ex vivo* HeLa cell screen, except serial compound dilùons in medium were prepared manually from 80 μM to ∼0.4 nM at 1.189× dilùon steps. Small dilùon steps rather than concentràon replicates were selected as it provides greater stàstical power from a set number of samples^71^. Final DMSO concentràon was 0.08% in all samples, and each plate included at least 12 control wells with 0.08% DMSO. Cytoplasmic FUS-GFP condensate number and nuclear/cytoplasm partition of FUS-GFP were analysed using custom macros in Fiji. Nuclei were identified by intensity thresholding of DNA images labelled with Hoechst following a 5 px Gaussian blur. Cytoplasmic FUS-GFP condensates were identified by intensity thresholding of the FUS-GFP images following a 10 px weight 0.9 unsharp filter masked by the thresholded nuclei. The rào of the number of cytoplasmic FUS-GFP condensates to that of nuclei was taken as cytoplasmic FUS-GFP condensates per cell per field of view, and *ρ*, the rào of partition of FUS-GFP to the nucleus and the cytoplasm, was derived from *α* = *ν*_*n*_⁄*ν*_*t*_, the rào of nuclear to total green signal per field of view, where *ρ* = *α*⁄(1 − *α*). These data were log transformed and fitted to a Rodbard sigmoidal curve^72^ to determine EC_50_. Six fields of view were captured and analysed per condìon.

The series of lipoamide analogues including lipoamide and lipoic acid were newly synthesized by Wuxi AppTec and provide through Dewpoint Therapeùcs. To assess those dose response effect, the HeLa BAC cells of FUS-GFP were seeded in 384-well plates (4000 cell per well) 24 h prior treatment, pre-treated with the compounds in a half-log dilùon series (from 30 µM to 3 nM: seven concentràons) using an Echo 650, and followed by a 1 h treatment with 1.5 mM potassium arsenate before fixàon with 4% formaldehyde for 15 min, permeabilizàon with 0.1% Triton X-100 for 10 min, and counter-staining with Hoechst and cell mask blue as described above. Imaging was performed using an Opera Phenix (PerkinElmer), 20×, 9 FOV, binning 2, and using Harmony 4.9 software to determine cytoplasmic FUS-GFP condensates number as well as cytoplasmic and nuclear FUS-GFP intensìes to calculate nuclear to cytoplasmic rào of FUS-GFP intensìes. EC_50_ was calculated either using CDD Vault curve fiãng or Harmony 4.9 software.

### *In vitro* protein condensation, solidification, and oxidation assays

For the condensàon assay at different KCl concentràons, FUS-GFP proteins in storage buffer was diluted with 20 mM HEPES pH 7.25 containing DMSO and lipoamide to give 20 µl of indicated concentràons of the protein and KCl, 0.3 mM DTT, and 300 µM lipoamide (0.3% DMSO), and placed on a 384-well plate (781096, Greiner). Condensates were imaged on a Nikon TiE inverted microscope with a Nikon Apo 60× NA 1.2 water immersion objective using a Yokogawa CSU-X1 spinning disk head and an Andor iXon EM+ DU-897 EMCCD camera

The assay to determine dilute phase concentràons at different temperature was performed with a newly established technique, which will be reported in detail elsewhere. In brief, the technique is based on mass and volume conservàon and defined reaction volumes. We can use this method to determine accurate values for both dilute and condensed branch protein concentràons. Here, FUS-GFP phase separàon was induced for a protein concentràon titràon in water-in-oil emulsions in a buffer containing 25 mM Tris-HCl pH 7.4, 150 mM KCl, 1 mM DTT, and the indicated concentràons of lipoamide (or DMSO as control) and imaged with a CSU-W1 (Yokogawa) spinning disk confocal system at an IX83 microscope with a 40x UPlanSApo 0.95 NA air objective, controlled via CellSens (Olympus). The dilute phase protein concentràon was derived from a linear fit to the volume fractions of FUS-GFP condensed phase versus the total concentràons of FUS-GFP. Temperature was controlled using a custom-made stage^73^.

For solidificàon assays, FUS-GFP in storage buffer was diluted in 50 mM Tris-HCl pH 7.4, 1 mM DTT to give 10 μM protein, 50 mM Tris-HCl pH 7.4, 1 mM DTT, 50 mM KCl in a 20 μl volume in non-binding clear bottom 384 well plates (781906, Greiner). Compounds, or an equal volume of DMSO, were then added for a final compound concentràon of 30 μM and 0.3% DMSO. ‘Aging’ to cause fibre formàon was induced by horizontal shaking at 800 rpm at room temperature, as previously described^12^. Fibre and condensate formàon were analysed by widefield epifluorescence using a DeltaVision Elite microscope (GE Healthcare Life Sciences) with a Plan ApoN 60× NA 1.4 oil immersion objective (Olympus) and an sCMOS camera (PCO). Fluorescence recovery after photobleaching (FRAP) of FUS-GFP condensates and fibres was performed on a Nikon TiE inverted microscope with a Nikon Apo 100× NA 1.49 oil immersion objective using a Yokogawa CSU-X1 spinning disc head and an Andor iXon EM+ DU-897 EMCCD camera. 10×10 px regions were bleached for 50 ns with a 6 mW 405 nm laser using an Andor FRAPPA beam delivery unit then imaged for 5 min at 5 Hz. Recovery curves were derived using scripts in Fiji.

Oxidàon of SFPQ was detected by change in mobility in SDS-PAGE without reducing agents. 10 µM of untagged SFPQ in buffer (20 mM HEPES pH 7.25 and 150 mM KCl) was incubated with H_2_O_2_ at RT for 30 min before subjecting to SDS-PAGE. For condensàon assays of individual proteins with H_2_O_2_, condensates of SFPQ-GFP and FUS-GFP were induced in buffer (20 mM HEPES pH 7.25 and 75 mM KCl). The assays for dissolùon and revival of FUS-SNAP condensates were performed in buffer (20 mM HEPES pH 7.25 and 150 mM KCl). FUS-SNAP was labelled with SNAP-Surface Alexa Fluor 546 (New England Biolabs), and protein mixtures were oxidized with H_2_O_2_ at RT for 1 h before image acquisìon. Proteins were imaged similarly to FUS-GFP condensates above.

### Controlled droplet fusion using optical tweezers

Liquidity of FUS protein condensates was assessed by controlled fusion experiments using dual-trap optical tweezers, as detailed previously^12,34^. In short, for each independent fusion event, two FUS protein droplets in the presence of 300 µM lipoamide or equivalent amount of DMSO (0.3%) as the control were trapped in each optical trap and brought into contact to inìate droplet coalescence. Fusion relaxàon times were accurately recorded as changes to the laser signal as condensate material flows into the space between the two optical traps during coalescence. The laser signal was recorded at 1 kHz, smoothed at 100 Hz and used to extract the characteristic relaxàon time. After fusion was complete –as indicated by a stable laser signal– the fused droplet was discarded, and two new droplets were captured for quantifying an independent fusion event.

### *Ex vivo* DNA cut assays

UV micro-irradiàon was performed as previously described^12,41^. Briefly, iPSCs expressing wild-type FUS-GFP were stressed by addìon of 1 mM arsenate for 1 h, then treated with lipoamide or an equal volume of DMSO for 1 h. A single point in the nucleus was subject to 3 UV pulses as described for FRAP, but at 10% laser power. GFP fluorescence was imaged at 1 Hz, and intensity of response was analysed on Fiji. iPSC-MNs expressing FUS P525L-GFP were pre-treated with 20 µM Lipoamide for 24 h before laser irradiàon. The UV laser cutter setup ùlized a 355-nm UV-A laser with a pulse length of <350 ps. A Zeiss alpha Plan-Fluar 100× 1.45 oil immersion objective was used, and 12 laser shots in 0.5 µm-steps were administered over a 12 µm linear cut.

### NMR for FUS-lipoamide interaction

Untagged FUS low complexity domain (residues 1 to 163) was expressed, purified, and analysed using _1_H-^15^N heteronuclear single quantum coherence NMR and sample condìons previously described^74^ in the presence of 500 μM lipoamide or equivalent DMSO solvent control (1%).

### NMR for lipoamide concentrations

#### Synthesis/validation of [15N]-labelled lipoamide

[15N]-racemic (±) and (*R*)-(+)-lipoamide were synthesized from racemic and (*R*)-(+)-lipoic acid, respectively, by activàng the carboxylic acid using N-hydroxysuccinimide and (1-ethyl-3-(3-dimethylaminopropyl) carbodiimide hydrochloride. The NHS derivàve was reacted with ^15^NH_4_Cl to incorporate the [15N]-labelling. Full details of the synthesis and subsequent biophysical validàon are included in Supplementary Informàon.

### NMR based detection and quantification of [15N]-labelled lipoamide

_1_H detected ^15^N edited ^1^H sensìvity enhanced HSQC NMR ((^15^N)^1^H) spectra were acquired on a 14.1 T Varian Inova spectrometer equipped with a 5 mm z-axis gradient triple resonance room temperature probe. The free induction decay was recorded for an acquisìon time of 0.0624 s and a sweep width of 8 kHz recorded over 1000 points and a recovery delay of 1 s. Typically, 10000 transients were collected giving a total experiment time of 3 h 1 min. The J coupling between the amide protons and the ^15^N in H_2_O samples was determined to be 88 Hz, and so the transfer times of 1/4 J in the INEPT portions of the pulse sequence were set to 2.6 ms. With these seãngs, ^15^N ammonia or ammonium ions would not be detectable. Chemical modificàon of [15N]-lipoamide (including covalent attachment to an apoenzyme) would give a substantial change in the (^15^N)^1^H NMR spectrum. Similarly, dissolùon of lipoamide in a phospholipid membrane would give substantial peak broadening in the cell samples. We observed neither, consistent with freely diffusing lipoamide.

### Optimisation of the NMR measurement conditions of [15N]-labelled lipoamide

Solvent, pH and temperature sensìvity of the primary amide proton chemical shifts were determined using dummy samples assembled from the appropriate solvent and added compounds.

Integrated NMR signal intensity is proportional to concentràon if provided condìons (temperature and pH) are identical^75^. Chemical exchange^76^, expected as the amide protons in lipoamide should be labile in water, must also be accounted for. To select appropriate condìons, we determined temperature (Figure S2F) and pH (Figure S2G) sensìvity of the amide proton signal of 1 mM [15N]-lipoamide in cell medium. Both amide protons showed chemical exchange under high temperature, high pH condìons, with the trans-amide proton affected weakly (Figure S2F and S2G). We then assessed degradàon of the trans-amide proton over 10 h (Figure S2H). At 37°C, but not 10°C, the signal intensity decayed slowly, suggesting slow hydrolysis to form ammonia. We concluded that at 10°C and below pH 8.6 the integrated signal from the trans-amide proton resonance is a good measure of [15N]-lipoamide concentràon.

### Quantification of [15N]-lipoamide cellular uptake

#### Sample preparation workflow

HeLa cells expressing FUS-GFP were grown in 6 well plates to 10^6^ cells/well in DMEM supplemented with 10% FCS. To simultaneously stress and treat cells, the medium was replaced with 0.6 ml medium supplemented with potassium arsenate and 100 μM [15N]-racemic (±) or (*R*)-(+)-lipoamide for 1 h at 37°C. High concentràons of compound were used to maximise the signal. The medium was then removed and retained (medium sample), the cells washed with ∼2 ml PBS, then the cells removed by trypsinisàon: addìon of 0.3 ml TrypLE Express (12604013, Thermo Fisher Scientific) and incubàon at 37°C for 5 min, then addìon of 0.3 ml medium to quench the trypsin. The resuspended cells were retained (cell sample). All samples were frozen at −80°C. Wells were prepared for all combinàons of no compound (1% DMSO control), [15N]-(±)-lipoamide or [15N]-(*R*)-(+)-lipoamide, with or without potassium arsenate and with or without cells.

#### Calculation of cell volume and uptake

The concentràons of [15N]-lipoamide inside (C_cell_) and outside (C_out_) the cells were calculated from measurements of signal intensity *S* of the trans-amide proton of lipoamide acquired in the absence (−*cells*, sample i, Figure S2A) and presence (+*cells*, sample ii, Figure S2A) of HeLa cells, using the following equàons (full derivàon included in Supplementary Informàon):

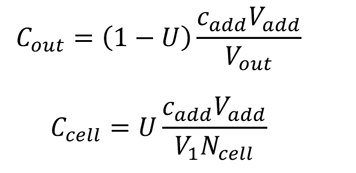

where *N*_&’((_ = 10^6^, *c*_$%%_ = 100 μM and *V*_$%%_ (added volume) = 600 μl, *V*_)_ = 4.19 x 10^-15^ m^3^ (approximàng HeLa cells as spheres of radius 10^-5^ m) and U represents measured fractional uptake as given by:

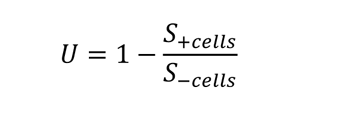

### *In vitro* partitioning of [15N]-(±)-lipoamide in FUS-GFP condensate phase

#### Sample preparation workflow

Phase separàon of FUS-GFP, at room temperature, was achieved by dilùng 12.5 μl of protein stock at 170 μM concentràon (in salty HEPES buffer – 50 mM HEPES, 500 mM KCl and 5% glycerol, at pH 7.25, DTT 1 mM) with 247 μl of salt-free buffer containing [15N]-(±)-lipoamide (50 mM HEPES, 5% D_2_O, 105 μM [15N]-(±)-lipoamide, 1.05% DMSO, at pH 8), resulting into samples of 260 μl with 8 μM FUS-GFP, 100 μM [15N]-(±)-lipoamide and 25 mM KCl.

The sample was centrifuged for 10 min at 4000 × *g* and room temperature and the supernatant kept for NMR analysis. The remaining supernatant was carefully pipetted out, without disturbing the condensate pellet, and discarded. Perpendicular view photographs of the pellet were taken. Finally, the condensate pellet was resuspended in 260 μl of buffer with the same buffer condìons as the phase separated sample (50 mM HEPES, 25 mM KCl, pH ∼7.4).

Resuspended condensate or supernatant were loaded in D_2_O-matched 5 mm Shigemi tubes and analysed by (^15^N)^1^H NMR. To achieve adequate signal to noise, the resuspended condensate was scanned for 20 h, while the supernatant was scanned for 4 h. The signal factor (intensity rào between the dilute and resuspended condensate samples) was adjusted accounting for differences in sample volume and number of scans.

#### Calculation of condensate phase volume

The volume of condensate was calculated from perpendicular photographs (see Fig. S3C), from the pellet radius (a) and inner radius of the semi-spherical bottom of the microcentrifuge tube (r):

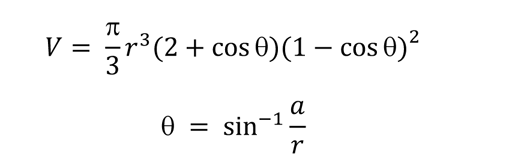

*0* = subtended angle, as showed in Fig. S3C.

Calculàon of partition coefficient

The rào between the concentràon of lipoamide in the condensate and dilute phases (partition coefficient - PC) was calculated using:

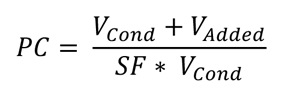

where *V*_/!0%_ is the volume of condensate phase, *V*_1%%’%_ is the volume added to resuspend the condensate phase (260 μl) and SF is the signal intensity rào between the dilute phase and the resuspended condensate measured by NMR.

The concentràons of lipoamide in the condensate ([L]_Cond_) and dilute ([L]_Dil_) phases were calculated as:

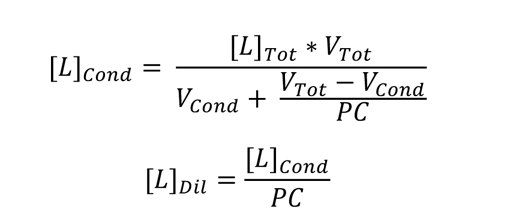

Where [L]_Tot_ is the total concentràon of [15N]-(±)-lipoamide and V_Tot_ is the total volume of the phase separated sample. The fraction (%) of [15N]-(±)-lipoamide signal in the condensate phase (see Fig. S3D) was calculated as:

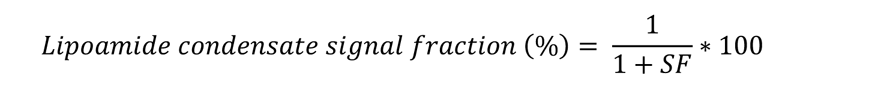

The full derivàon of these expressions can be found in the Supplementary Informàon.

#### Crosslinking coupled to Mass Spectrometry (XL-MS)

FUS condensates were processed and analyzed essentially as described previously^77^. In short, reconstituted droplets of lysine-rich FUS K12 or FUS G156E were generated by low salt (80 mM KCl) and crosslinked by addition of H12/D12 DSS (Creative Molecules) in the presence or absence of lipoamide for 30 min at 37°C, shaking at 600 rpm. Protein samples were quenched by addition of ammonium bicarbonate to a final concentration of 50 mM and directly evaporated to drynes. The dried protein samples were denatured in 8 M Urea, reduced by addition of 2.5 mM TCEP at 37°C for 30 min and subsequently alkylated using 5 mM Iodoacetamide at RT for 30 min in the dark. Samples were digested by addition of 2% (w/w) trypsin (Promega) over night at 37°C after adding 50 mM ammonium hydrogen carbonate to a final concentration of 1 M urea. Digested peptides were separated from the solution and retained by a C18 solid phase extraction system (SepPak Vac 1cc tC18 (50 mg cartridges, Waters) and eluted in 50% ACN, 0.1% FA. Dried peptides were reconstituted in 30% ACN, 0.1% TFA and then separated by size exclusion chromatography on a Superdex 30 increase 3.2/300 (GE Life Science) to enrich for crosslinked peptides. Peptides were subsequently separated on a PepMap C18 2 µM, 50 µM x 150 mm (Thermo Fisher Scientific) using a gradient of 5 to 35% ACN for 45 min. MS measurement was performed on an Orbitrap Fusion Tribrid mass spectrometer (Thermo Fisher Scientific) in data dependent acquisition mode with a cycle time of 3 s. The full scan was done in the Orbitrap with a resolution of 120000, a scan range of 400-1500 m/z, AGC Target 2.0e5 and injection time of 50 ms. Monoisotopic precursor selection and dynamic exclusion was used for precursor selection. Only precursor charge states of 3-8 were selected for fragmentation by collision-induced dissociation (CID) using 35% activation energy. MS2 was carried out in the ion trap in normal scan range mode, AGC target 1.0e4 and injection time of 35 ms. Data were searched using *xQuest* in ion-tag mode. Carbamidomethylation (+57.021 Da) was used as a static modification for cysteine. Crosslinks were quantified relative to the condition containing no lipoamide.

#### Transfection

Transfection for gene deplèon was performed with Lipofectamine2000 (Thermo Fisher Scientific) and esiRNA oligos targèng human genes (Eupheria Biotech), as listed in Table S1. esiRNA targèng Renilla luciferase was used for a negàve control. The medium was replaced 5 h after transfection, and the cells were cultured for 3 days before analysis.

#### Immunocytochemistry of cultured cells

HeLa cells were fixed with 4% paraformaldehyde (PFA) in PBS at room temperature for 15 min, then washed with PBS containing 30 mM glycine. After permeabilizàon with 0.1% Triton X-100 in PBS at 4°C and the following wash with glycine-containing PBS, the cells were blocked with 0.2% fish skin gelàn (Sigma) in PBS (blocking buffer) at room temperature for 20 min, incubated with primary antibodies in blocking buffer overnight at 4°C, washed with blocking buffer, and incubated with secondary antibodies and DAPI in blocking buffer at room temperature for 1 h. After washed with blocking buffer, the samples were stored in PBS until imaging. For detecting endogenous SFPQ, cells were fixed with cold methanol on ice for 10 min, and blocked with blocking buffer at room temperature for 20 min before treated with primary antibodies. Samples were imaged on Zeiss LSM 700 or 880 confocal microscopes with a 40× NA 1.2 water objective (Zeiss). Segmentàon of nuclei, the cytoplasm, stress granules, and mitochondria, and measurement of fluorescence intensìes at each segment were performed using CellProfiler. The data were then processed using KNIME to calculate number of stress granules per cell, nuclear-to-cytoplasmic intensity rào of stress granule proteins, intensity rào of the click-crosslink lipoamide analogue (i) at stress granules, mitochondria, and nuclei over the cytoplasm (excluding stress granules and mitochondria) or (ii) without crosslinking over with crosslinking, and percent of cells that have more than 2 stress granules.

iPSC-MNs were fixed for 15 min at room temperature in 4% PFA in PBS. Permeabilizàon and blocking was performed simultaneously using 0.1% Triton X-100, 1% BSA and 10% FBS in PBS at room temperature for 45 min. Subsequently, the primary antibodies were applied overnight at 4 °C in 0.1% BSA in PBS. The cells were washed with 0.1% BSA in PBS and incubated with secondary antibodies for 1 h at room temperature. Finally, the cells were washed three times with 0.1% BSA in PBS-T (0.005% Tween-20), including Hoechst or DAPI in the second washing step. Neurofilament H (NF-H) was used for a marker of MNs. Samples were imaged on either a CellVoyager CV7000 automated spinning disc confocal microscope (Yokogawa) with a 40× NA 1.3 water objective or a Zeiss LSM880 laser scanning confocal microscope.

The following primary antibodies were used: rabbit anti-G3BP1 (PA5-29455, Thermo Fisher Scientific); mouse anti-Tom20 (F-10, Santa Cruz); mouse anti-SPFQ (C23, MBL); mouse anti-SRSF1 (103, Invitrogen); rabbit anti-TDP-43 (80002-1-RR, Proteintech); mouse anti-Neurofilament H (SMI-32, Millipore); mouse anti-ý3 Tubulin (T5076, Sigma-Aldrich). The secondary antibodies used are as follows: Alexa Fluor 488-conjugated anti-mouse, Alexa Fluor 594-conjugated anti-rabbit, anti-mouse, and Alexa Fluor 647-conjugated anti-rabbit and anti-mouse (Thermo Fisher Scientific).

#### UV cross linking and click reaction

HeLa cells were treated with 3 mM arsenate for 1 h, followed by 30 µM of the click-crosslink lipoamide analog for 30 min in the presence of arsenate. Then the cells were irradiated with a 305 nm light-emiãng diode for 10 sec for cross linking just before fixàon with 4% PFA in PBS at room temperature. The fixed cells were subjected to immunostaining as described above. After staining, the cells were subjected to click reaction with 2 µM AF594-Picolyl-Azide (CLK-1296-1, Jena Bioscience) in buffer containing 100 mM HEPES pH 7.25, 5 mM L-ascorbic acid, 0.5 mM THPTA, and 0.1 mM CuSO_4_ at 37°C for 40 min. Cells were then washed three times with 0.1% Triton X-100 in PBS to remove free dye. Imaging was performed on CSU-W1 (Yokogawa) spinning disk confocal system on an IX83 microscope (Olympus) with a 100× UPlanSApo 1.4 NA oil objective (Olympus).

#### Treatment with L-azidohomoalanine (AHA)

Wilde-type HeLa cells were firstly washed with and cultured in methionine (Met)-free medium (Gibco 21013-24) supplemented with 10% FBS for 1 h. Then the medium was replaced with complete medium or Met-free medium supplemented with 1 mM of Met (Sigma M9625) or AHA (Invitrogen C10102) for 2 h before the cells were stressed with 1 mM arsenate for 1 h. After fixàon with 4% PFA in PBS, the cells were subjected to immunostaining to label G3BP1.

#### Time-lapse imaging

Time-lapse imaging was performed at 37°C with 5% CO_2_. iPSC-derived MNs were treated with 20 µM lipoamide or 0.02% DMSO (control) for 1 h and then treated with 20 µM arsenite just before image acquisìon. Maximum projection images were generated, and number of FUS-GFP foci was quantified by Fiji.

#### Axonal transport assays

AH-ALS1-F58 iPS MNs expressing P525L FUS were treated with 2 μM compound or an equal volume of DMSO for 3 days. Longer incubàon was selected to ensure penetràon and action of compounds along the length of the axon channel. 2 μM was selected as the highest concentràon where there were no toxic effects on this iPS line (assessed qualitàvely). Analysis of axonal transport of lysosomes were performed as previously described^41^. Briefly, lysosomes were labelled by addìon of 50 nM lysotracker red (Thermo Fisher Scirntific) and imaged using a Leica DMI6000 inverted microscope with a 100× NA 1.46 oil immersion objective and an Andor iXON 897 EMCCD camera in an incubator chamber (37°C, 5% CO2) at 3 Hz for 120 s at either the proximal or distal end of the silicone channels harbouring the axons. Kymographs were generated on Fiji. Particle tracking was used to identify proportion of particles moving faster than 0.2 μm/s for five videomicrographs. Each video includes a variable populàon of non-moving background particles, therefore, for each biological replicate, data were normalised to the mean proportion of moving lysosomes (>0.2 µm/s) at either MFC site (proximal and distal) in the DMSO (solvent control)-treated FUS P525L samples in Fig. 7D.

#### Protein aggregation in *C. elegans*

The effect of lipoic acid on stress granule protein aggregàon *in vivo* was analysed using a *C. elegans* model for stress granule formàon and aggregàon. As previously described^44,78^, fluorescent-tagged PAB-1 forms abundant stress granules and large solid aggregates during aging or upon chronic stress. RHO-1 also aggregates during aging, but is not an RNA binding or stress granule protein. Two lines were used: Fluorescently tagged PAB-1 (DCD214: N2; *uqIs24[pmyo-2::tagrfp::pab1gene*]) and RHO-1 (DCD13: N2; *uqIs9[pmyo-2::rho-1::tagrfp+ptph-1::gfp*]). Each were analysed as below, except DCD13 were maintained at 20°C.

The animals were exposed to lipoic acid in liquid culture in a 96 well plate starting from larval stage L4 in a total volume of 50 μl S-Complete per well (100 mM NaCl, 50 mM Potassium phosphate pH 6, 10 mM potassium citrate, 3 mM MgSO_4_, 3 mM CaCl_2_, 5 μg/mL cholesterol, 50 μM ethylenediaminetetraacèc acid (EDTA), 25 μM FeSO_4_, 10 μM MnCl_2_, 10 μM ZnSO_4_, 1 μM CuSO_4_) supplemented with heat-killed OP50 and 50 μg/ml carbenicillin. Per experiment, a minimum of nine wells each with 13 animals were treated with R-(+) or S-(−)-lipoic acid or an equivalent volume of DMSO.

48 h after switching the L4s from 20°C to 25°C (day 2 of adulthood) extensive aggregàon of fluorescently tagged PAB-1 and RHO-1 occurs in the pharyngeal muscles. After immobilizàon with 2 mM levamisole aggregàon was scored using a fluorescent stereo microscope (Leica M165 FC, Plan Apo 2.0× objective). For PAB-1, aggregates occurred primarily in the terminal bulb of the pharynx, and aggregàon was scored by the number of aggregates (>10 per animal). For RHO-1, aggregates were scored in the isthmus of the pharynx and aggregàon was scored as high (>50% of the isthmus), medium (<50%) or low (no aggregàon). High-magnificàon images were acquired with a Leica SP8 confocal microscope with a HC Plan Apo CS2 63× NA 1.40 oil objective using a Leica HyD hybrid detector. tagRFP::PAB-1 was detected using 555 nm as excitàon and an emission range from 565-650 nm. Representàve confocal images are displayed as maximum z stack projection.

#### D. melanogaster ALS models

All fly stocks were maintained on standard cornmeal at 25°C in light/dark controlled incubator. *w^1118^*, *UAS-eGFP*, *D42-GAL4*, and *OK6-Gal4* flies were obtained from Bloomington Drosophila Stock Center. *UAS-FUS WT*, *UAS-FUS P525L*, and *UAS-FUS R521C* flies were previously described^45,79^. *UAS-TDP-43 WT* and *UAS-TDP-43 M337V* flies were provided by J. Paul Taylor^80^.

Tissue-specific expression of the human genes was performed with the Gal4/UAS-system^81^. Climbing assays were performed as previously described^79^. Briefly, flies expressing eGFP, human FUS, or TDP-43 were grown in the presence or absence of Lipoic Acid (430 µM, ethanol as the vehicle control) or Lipoamide (430 µM, DMSO as the vehicle control), then anesthèsed, placed into vials and allowed to acclimàse for 15 min in new vials. Feeding these compounds did not show obvious lethality or toxicity at this concentràon. For each fly genotype, the vial was knocked three times on the base on a bench and counted the flies climbing up the vial walls. The percentage of flies that climbed 4 cm in 30 s was recorded. TDP-43-expressing flies were raised at 18°C to suppress lethality.

For immunohistochemistry of neuromuscular junctions, parent flies were crossed on food supplemented with DMSO or lipoamide, and the offspring were raised on the same food. Wandering third instar larvae were dissected and subjected to immunostaining as described previously^82^. Briefly, the dissected larvae were fixed with 4% PFA in PBS at room temperature for 20 min, then washed with PBS. After removing unnecessary tissues, the samples were blocked with 0.2% fish skin gelàn (Sigma) and 0.1% Triton X-100 in PBS (blocking buffer) at room temperature for 1 h, incubated with anti-HRP-Cy3 (1:200, Jackson Immunoresearch) in blocking buffer overnight at 4°C, washed with 0.2% Triton X-100 in PBS (PBT), and incubated with Alexa Fluor 488 Phalloidin (1:5000, Thermo Fisher Scientific) at room temperature for 2 h to visualize muscles. The samples were then washed with PBT and mounted with 70% glycerol in PBS. Synaptic boutons of muscle 4 in abdominal segments 2, 3, and 4 (A2–A4) were imaged using Zeiss LSM 700 or 880 confocal microscopes with a 40× NA 1.2 water objective (Zeiss). Numbers of synaptic boutons and satellite boutons were counted manually.

#### Quantitative RT-PCR (qPCR)

qPCR was performed with primers targèng GAPDH (control) and full-length STMN2, as described previously^83^.

#### Thermal proteome profiling (TPP)

Thermal proteome profiling was performed as described previously^31^. In brief, two 150 mm dishes of HeLa cells (∼6 million cells per dish) were treated with 0.1% (v/v) DMSO (control) or 100 µM lipoamide for 1 h. At the end of incubàon one lipoamide-and one DMSO-treated dishes of HeLa cell were stressed with 1 mM arsenate for 1 h. The second set of cells served as the control (treatment with water, vehicle in which arsenate was dissolved) for only lipoamide treatment and only DMSO treatment. All incubàons were performed at 37C° with 5% CO_2_. Following the incubàon, the cells were washed with PBS and trypsinized. The cells were collected by centrifugàon at 300 × *g* for 3 min. The cell pellet was re-suspended in PBS containing the appropriate treatment concentràons of the compounds (lipoamide, DMSO, and arsenate) at cell density of 4 × 10^6^ cells/ml. This cell suspension was split into 10 × 100 µl aliquots on a PCR plate, spun at 1000 × *g* for 3min, and finally 80 µl of supernatant (PBS) was subsequently removed. The cell aliquots were then heated to ten to different temperatures (37.0, 40.4, 44.0, 46.9, 49.8, 52.9, 55.5, 58.6, 62.0, and 66.3°C) for 3 min in a thermocycler (SureCycler 8800, Agilent) and left at room temperature for 3 min. Subsequently, the cells were lysed with 30 µl of lysis buffer (PBS containing protease inhibitors, 1.12% NP-40, 2.1 mM MgCl_2_, and phosphatase inhibitors), and the PCR plates containing the cell lysate was centrifuged at 1000 × *g* for 5 min to remove cell debris. Next, the heat-induced protein aggregates were removed from the cleared supernatant by passing it through a 0.45 µm 96-well filter plate (Millipore) at 500 × *g* for 5 min. Equal volumes of the flow-through and 2× sample buffer (180 mM Tris-HCl pH 6.8, 4% SDS, 20% glycerol, and 0.1 g bromophenol blue) were mixed and stored in –20°C until used for mass spectrometry sample preparàon. Protein digestion, peptide labelling and mass spectrometry-based proteomics were performed as previously described^32^.

#### Data analysis of TPP

Abundance and thermal stability scores for every protein was calculated as described previously^30,84^. Briefly, the ratio of the normalized tandem mass tag (TMT) reporter ion intensity in each treatment (only lipoamide, only arsenate, lipoamide and arsenate) and the control (only DMSO) was calculated for each temperature. The abundance score for each protein was calculated as the average log_2_ fold change (FC) at the two lowest temperatures:

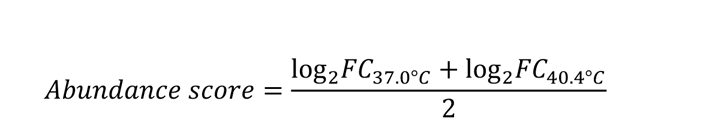

The thermal stability score for each protein was computed by subtracting the abundance score from the log_2_ fold changes of all temperatures, and then summing the resulting fold changes (requiring that there were at least ten data points to calculate this score):

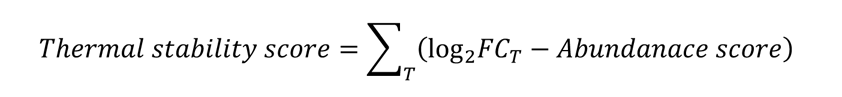

where *T* is the ten temperatures. Both the abundance and the thermal stability scores were transformed into a z-distribution by subtracting the mean and dividing by the standard deviation. The significance of the abundance and thermal stability scores was further assesses using *limma*^85^ (the two scores were weighted for the number of temperatures in which a protein was identified), followed by FDR estimation using the *fdrtool* package^86^. Proteins with |z-score| > 1.5 and with FDR < 0.05 were considered to be significant changes for the IDR analyses. Thermal stability scores indicated in the result section are those in cells treated with lipoamide and arsenate.

#### Bioinformatics

Posìons of amino acid sequences with disordered tendency were visualized using IUPred3 (hUps://iupred.elte.hu/). Length of IDRs in each protein was estimated using d2p2 database (hUps://d2p2.pro/)^87^. The IDR is defined as a region which is regarded as being disordered in more than 75% of all predictors in the database as well as with more than ten successive disordered amino acids. Then proportion of IDR(s) to the whole protein amino acid length was calculated. Enrichment of individual amino acids in IDRs were calculated using Composìon Profiler (hUp://**Error! Hyperlink reference not valid.** IDRs of all the proteins detected in TPP were used as a background. Posìve and negàve scores indicate enrichment and deplèon of each amino acid compared to the background, respectively.

#### Statistical analysis

Stàstical analyses were performed using the R stàstical software or GraphPad Prism. The stàstical details (the *p*-value, the number of samples, and the stàstical test used) are specified in the figure panels or legends. A *p*-value below 0.05 was considered stàstically significant.

#### Data availability

The mass spectrometry proteomics data have been deposited to the ProteomeXchange Consortium via the PRIDE^89^ partner repository with the dataset identifiers PXD039670 (for the crosslinking assay) and PXD039501 (for the TPP assay).

## Acknowledgements

We thank J.P. Taylor for providing fly strains; the following services and facilìes of the MPI-CBG for their support: fly team, LMF, PEPC, TDS, Scientific compùng facility, and OSCF; J. Robertson for his advice in the characterizàon of [15N]-labelled lipoamide; C. Möbius for helping on plate-based cell imaging; C. Höge for valuable comments on the manuscript; members in the Hyman lab and in the MPI-CBG for valuable discussion.

## Funding

Hiroyuki Uechi Uehara Memorial Foundation; JSPS Overseas Research Fellowships; The Osamu Hayaishi Memorial Scholarship for Study Abroad

Marcus Jahnel The Deutsche Forschungsgemeinschaft (DFG, German Research Foundation) under Germany’s Excellence Strategy – EXC-2068 – 390729961

Simon Alberti European Research Council (grant agreement number 725836)

Nicolas L. Fawzi Nàonal Institute of General Medical Sciences (NIGMS) of the Nàonal Institutes of Health (R01GM147677) and the Nàonal Science Foundàon (1845734)

Della C. David Deutsches Zentrum für Neurodegeneràve Erkrankungen (DZNE)

Andreas Hermann NOMIS foundation, an unrestricted grant by a family of a deceased ALS patient, the Stiftung zur Förderung der Hochschulmedizin in Dresden and the Hermann und Lilly Schilling-Stiftung für medizinische Forschung im Stifterverband.

Florian Stengel The German Research Foundation (STE 2517/1-1 and STE 2517/5-1)

Richard J. Wheeler Wellcome Trust Sir Henry Wellcome (211075/Z/18/Z) and Sir Henry Dale (103261/Z/13/Z) Fellowships

Satoshi Kishigami Ishizaka Memorial Foundation; Ezoe Memorial Foundation; Shigeta Foundation

BGD and AJB Next Generation Chemistry at the Rosalind Franklin Institute is funded by the EPSRC EP/V011359/1

## Supplemental figures

**Fig. S1.**
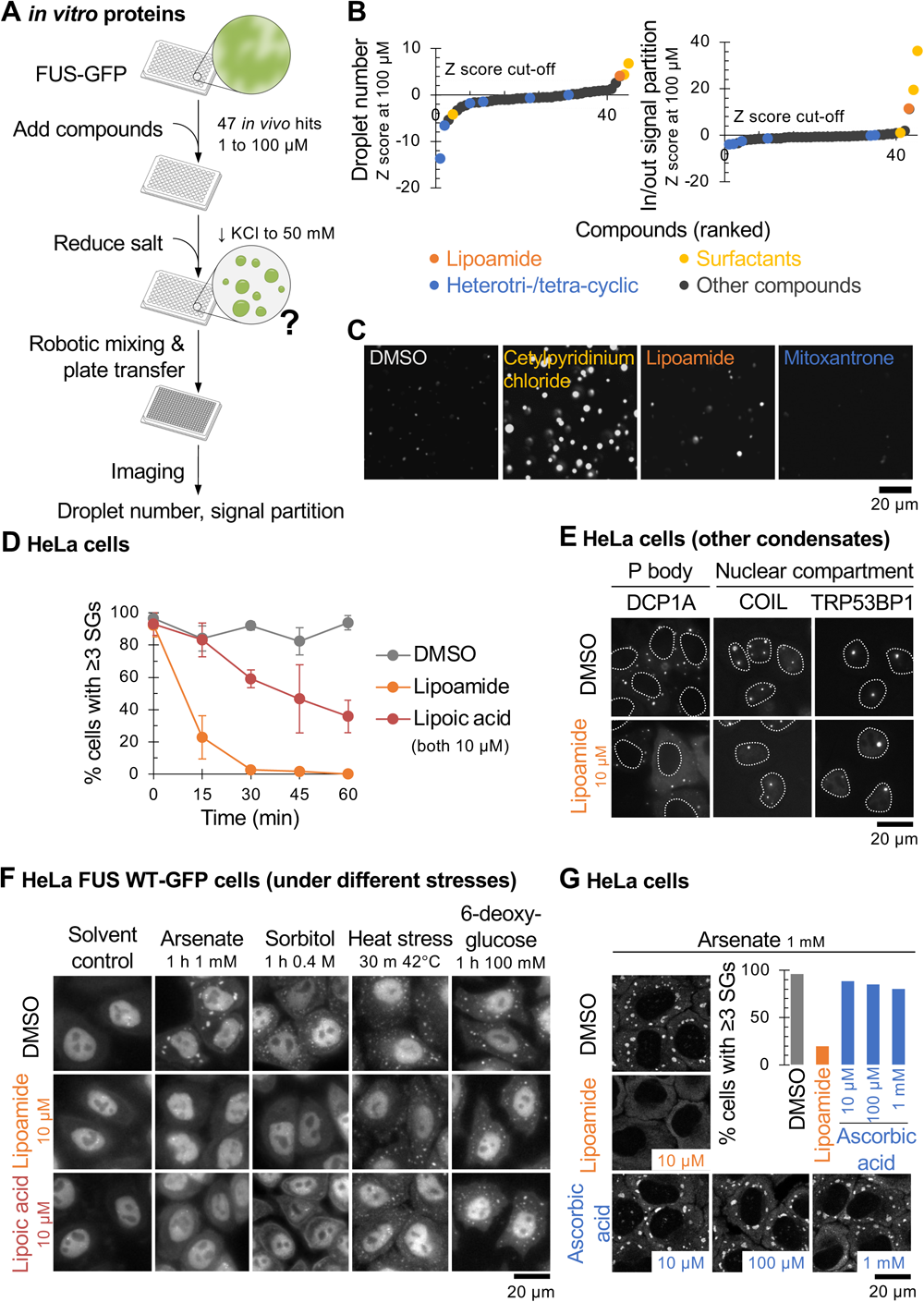
*In vitro* follow-up screening and lipoamide characterization in HeLa cells. **A,** Workflow for screening small molecules for effects on FUS condensàon of purified FUS-GFP *in vitro*. **B,** Ranked Z scores of change in condensate droplet number and signal partition into FUS-GFP droplets (formed under low salt condìons) where larger posìve or negàve values mean more compound effect. Scores were calculated at the maximum concentràon at which the compound solvent (DMSO) negàve control had no significant effect; 100 μM. Lipoamide, surfactant and heterotri-/tetracyclic compounds are indicated by data point colour. **C,** Appearance of the droplets with compound solvent (DMSO) negàve control or examples of compound classes: cetylpyridinium chloride (surfactant), lipoamide or mitoxantrone (heterotricyclic). Note the larger drops with cetylpyridinium chloride and lipoamide and the fewer smaller drops with mitoxantrone. **D,** Mean ± s.d. of percentage of HeLa cells with ≥ 3 G3BP1-posìve stress granules (SGs). Cells were treated with 1 mM arsenate for 1 h to induce SGs, followed by 10 µM lipoamide, lipoic acid, or 0.1% DMSO (control) in the presence of arsenate for indicated minutes. *n* = 52–248 cells from 3 independent experiments. **E,** Images of HeLa cells expressing GFP-tagged markers of other membrane-less organelle compartments subjected to 1 h treatment with 10 μM lipoamide (or DMSO control). Where unclear, the posìon of nuclei is indicated with a broken outline. Lipoamide does not disrupt P bodies (DCP1A), Cajal bodies (COIL), or DNA damage foci (TRP53BP1). **F,** Images of HeLa cells expressing FUS-GFP subjected to different stresses – arsenate, sorbitol (osmòc), heat, or 6-deoxyglucose (glycolysis) – with concurrent treatment with 10 μM lipoamide or lipoic acid. **G,** Representàve images of HeLa cells treated with 1 mM arsenate for 1 h, followed by 0.1% DMSO (control), 10 µM lipoamide, or indicated concentràons of ascorbic acid for 15 min. SGs were labelled with G3BP1.

**Fig. S2.**
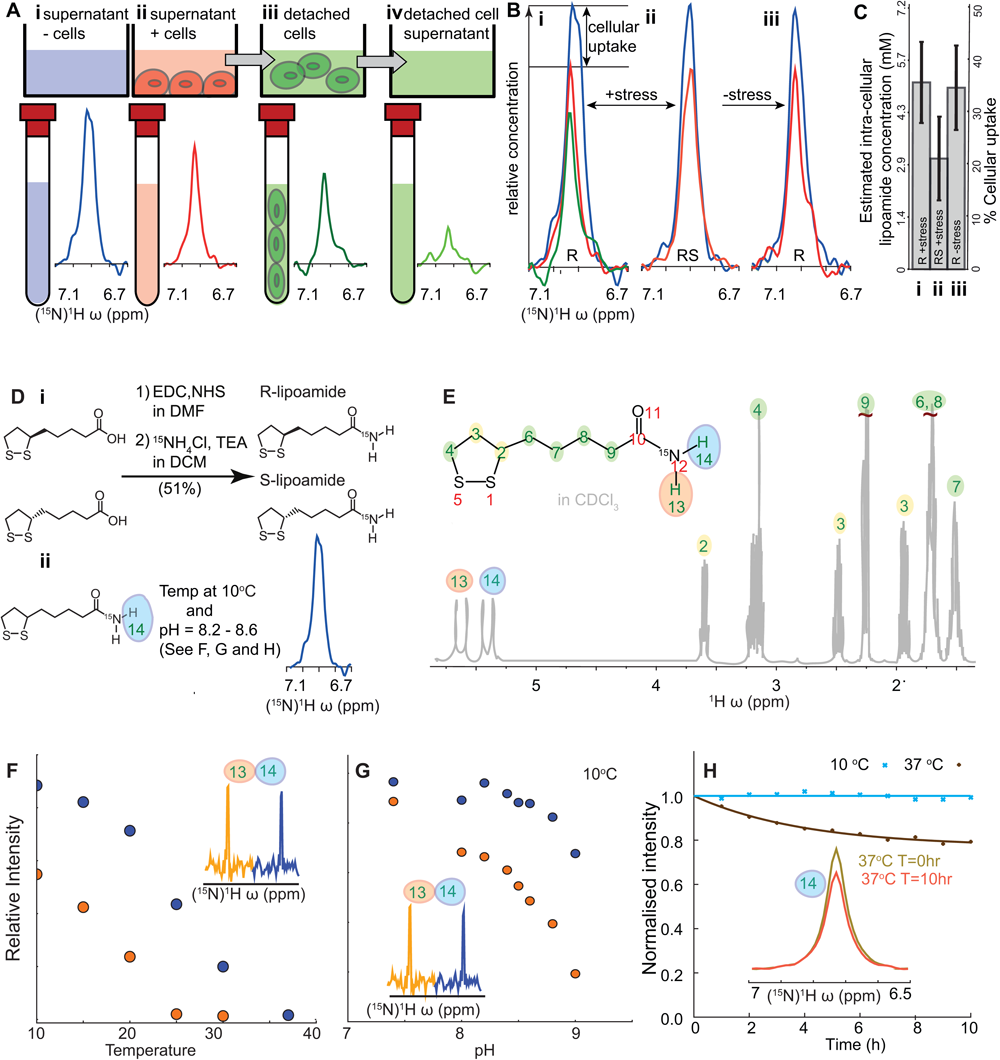
Tracking cellular uptake of [15N]-Lipoamide using NMR. **A,** Methodology for quantitàon of [15N]-lipoamide uptake by HeLa cells, using the trans-amide proton to measure [15N]-lipoamide concentràon (see F-H). Medium with 100 μM [15N]-lipoamide was incubated for 1 h in the absence or presence of HeLa cells. Following removal of medium, the cells were washed with medium (without arsenate) and detached using EDTA-trypsin. Solùon or cell pellet/in-cell NMR was used to determine [15N]-lipoamide concentràon. Example spectra for cells stressed with 3 mM arsenate and incubated with R-(+)-lipoamide are shown with the same y axis scale. **B,** Cellular uptake was determined by subtracting signal from medium incubated with cells (red) from signal from medium without cells (blue). This was carried out for all four combinàons of stressed (3 mM arsenate) or unstressed cells with [15N]-(*R*)-(+) or (±)-lipoamide. For stressed cells treated with [15N]-(*R*)-(+)-lipoamide the high signal intensity from the washed cell sample (green) is consistent with the large uptake from the medium calculated from the with (red) and without cell (blue) signal intensity. **C,** Quantitàon of B showing percentage uptake and calculated intracellular concentràon, assuming that lipoamide is uniformly distributed within cells (see Supplemental Methods). Uncertainty in measurement was approximately 30% and there was no significant difference in uptake between condìons. All measurements indicated substantial uptake of lipoamide and cellular concentràons >1 mM. **D,** Overview of synthesis of [15N]-lipoamide, highlighting the trans amide proton (14). **E,** ^1^H NMR spectrum of [15N]-lipoamide in CDCl_3_. Peaks can be unambiguously assigned to individual proton environments. F-H, Controls determining reliability of quantitàon of [15N]-lipoamide using the amide protons in ^15^N edited ^1^H NMR experiments. **F,** Dependency of the cis (13) and trans (14) amide proton signal on temperature, at a constant pH of 8.3. Both resonances decreased with increasing temperature, indicàng local molecular dynamics and/or interactions with H_2_O on ms to µs timescale reduce the signal. Trans amide proton resonance approaches a plateau towards 10°C. **G,** Dependency of the cis and trans amide proton signal on pH, at a constant temperature of 10°C. Together, indicàng at 10°C and below pH 8.6 integrated signal intensity of the trans-amide proton of lipoamide in ^15^N edited ^1^H NMR experiments is a reliable proxy for concentràon. **H,** Signal intensity of the trans-amide proton of lipoamide, when dissolved in growth medium, decreased over time at 37°C but not at 10°C. At 10°C signal intensity is stable for >10 h experiments.

**Fig. S3.**
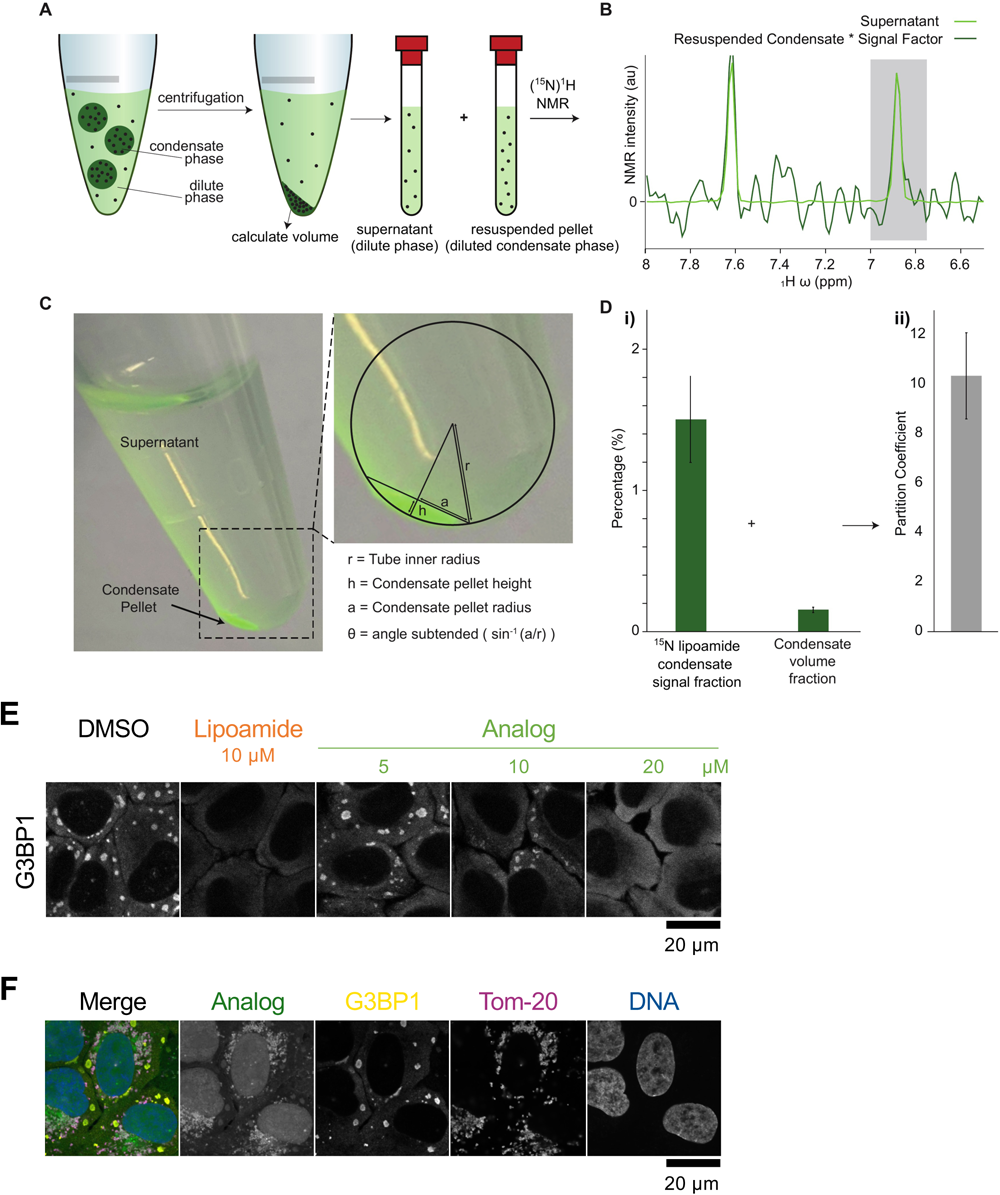
Experimental set up for portioning assays of lipoamide and its analogue. **A-D,** Methodology for determinàon of partition of [15N]-lipoamide into FUS condensates *in vitro.* (A) Schemàc showing the sample preparàon process. (B) Example of ^15^N edited ^1^H NMR signal around the ^15^N cis and trans amide protons for a dilute phase and condensate phase. Condensate phase spectrum is shown scaled by an experimentally determined signal factor, used in calculàon of the signal fraction. (C) Measurements for calculàon of condensate pellet volume from macro photographs of the sample within a microcentrifuge tube. (D) Mean ± s.d. of measured ^15^N edited ^1^H NMR signal fraction and condensate volume fraction (from 4 independent experiments) and calculated partition coefficient. Alternàve presentàon of the data in Fig. 2A. **E**, Representàve images of HeLa cells pre-treated with indicated concentràons of lipoamide or the click-crosslink lipoamide analog in Fig. 2B for 1 h followed by 1 mM of arsenate for addìonal 1 h in the presence of compounds. SGs were labelled with G3BP1. **F**, The images of HeLa cells treated with the analog and subjected to arsenate treatment and UV cross-linking from Fig. 2C, with a channel of Tom-20 as a mitochondrial marker.

**Fig. S4.**
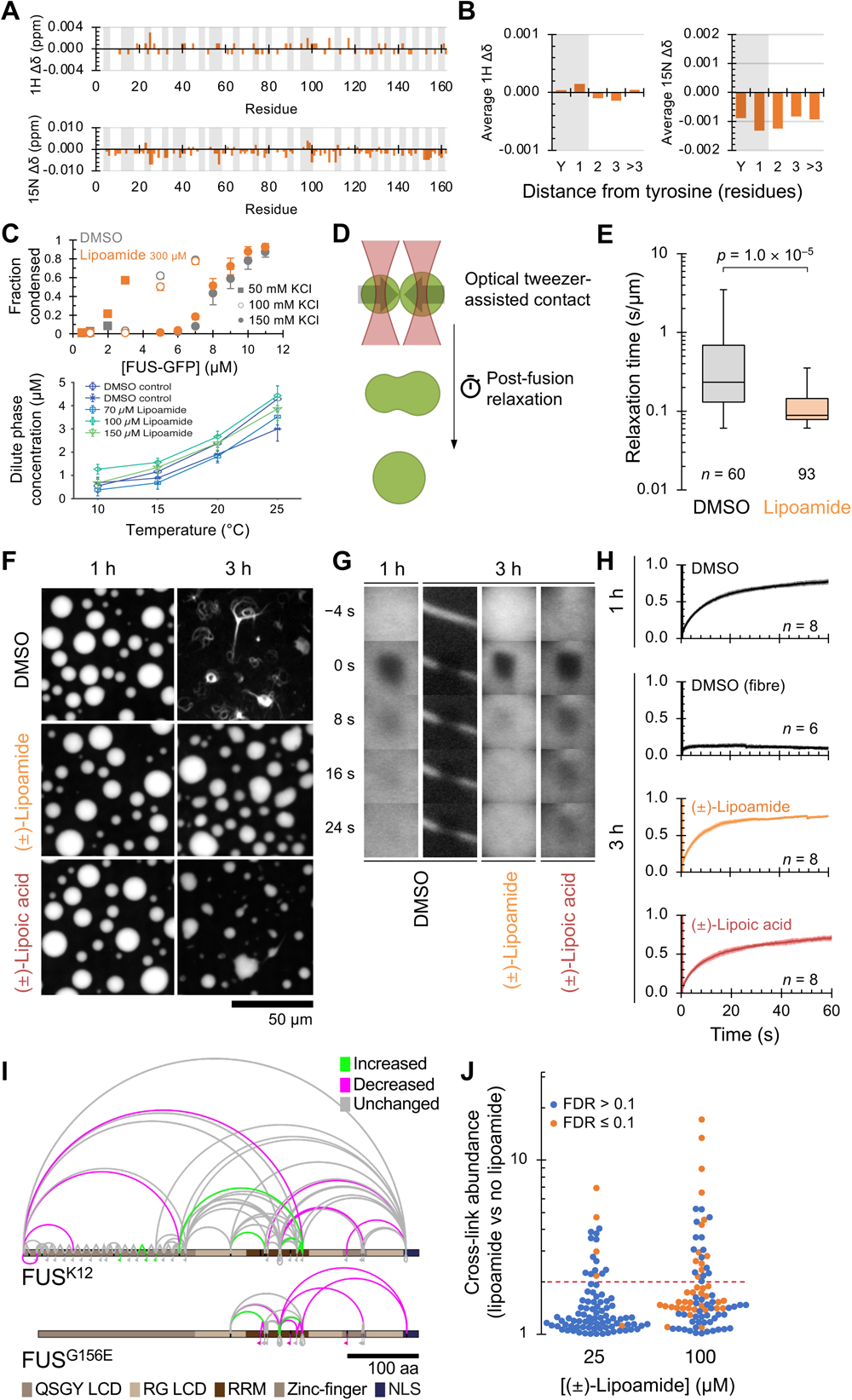
Lipoamide weakly increases liquidity of FUS condensates *in vitro*. **A,** NMR chemical shift deviàons per residue for the FUS N-terminal PLD (residues 1 to 163) with 500 μM lipoamide compared to the drug solvent control (1% DMSO). Light grey bars indicate tyrosine residues and residues neighbouring a tyrosine. **B,** Average ^1^H and ^15^N shifts across residues zero, one, two, three or more than three residues from a tyrosine in the presence of lipoamide. **C**, Top, mean ± s.d. of fractions of FUS proteins condensed at indicated salt (KCl) concentràons in the presence of 300 µM lipoamide or the DMSO control (0.3% v/v). *n* = 16 image fields. Bottom, dilute phase concentràons (equivalent to saturàon concentràons) of FUS-GFP at 150 mM KCl at different temperatures and lipoamide concentràons (errors are s.d.) **D**, Schemàc illustràng the quantitàon of condensate droplet liquidity using optical tweezers. Two droplets are brought into contact and begin to fuse: the time taken to relax to a single spherical droplet (once adjusted for the geometric mean radius as the characteristic droplet size) is a measure of the viscosity to surface tension rào of the droplet – a proxy of liquidity. **E**, Droplet size-corrected relaxàon times for droplet fusions with either 300 μM lipoamide (*n* = 93 independent fusion event) or equivalent DMSO solvent control (0.3%, *n* = 60). Box represents the 25^th^, 50^th^ and 75^th^ percentiles, whiskers represent 5^th^ and 95^th^ percentiles. *p* value by unpaired *t*-test. Lipoamide reduces fusion time, indicàng lower viscosity and/or greater surface tension. **F– H**, Effect of 30 μM lipoamide or lipoic acid on FUS G156E-GFP condensates ‘aging’, relàve to an equivalent DMSO solvent control (0.3%). Condensates were formed under 50 mM of KCl while shaking. (F) Representàve images after 1 and 3 h aging, showing fibre formàon in the DMSO sample in contrast to the lipoamide or lipoic acid samples. (G) Representàve fluorescence recovery after photobleaching (FRAP) time series of FUS condensates and fibres at corresponding time points. (H) Mean ± s.d. of relàve intensity of FUS-GFP FRAP in (G). Aged (3 h) condensates treated with lipoamide or lipoic acid maintain large FUS-GFP mobile fraction. Both compounds delay fibre formàon. **I**, Changes in intramolecular crosslinking due to lipoamide of FUS in *in vitro* low salt (80 mM KCl) condensates using the lysine-rich FUS K12 or FUS G156E. Significantly changed crosslinking sites with a change in intensity of more than two-fold and FDR ≤ 0.1; 3 independent experiments) are shown coloured in green (increased) or red (decreased). Other crosslinking sites are shown in grey. **J,** Dose-dependent effect of lipoamide on FUS K12, ploãng absolute change in crosslink intensity relàve to no lipoamide. Crosslinking sites with false discovery rate (FDR) > 0.1 are shown in blue, those with FDR ≤ 0.1 in orange (2 independent experiments). Two-fold change is indicated with a dashed red line.

**Fig. S5.**
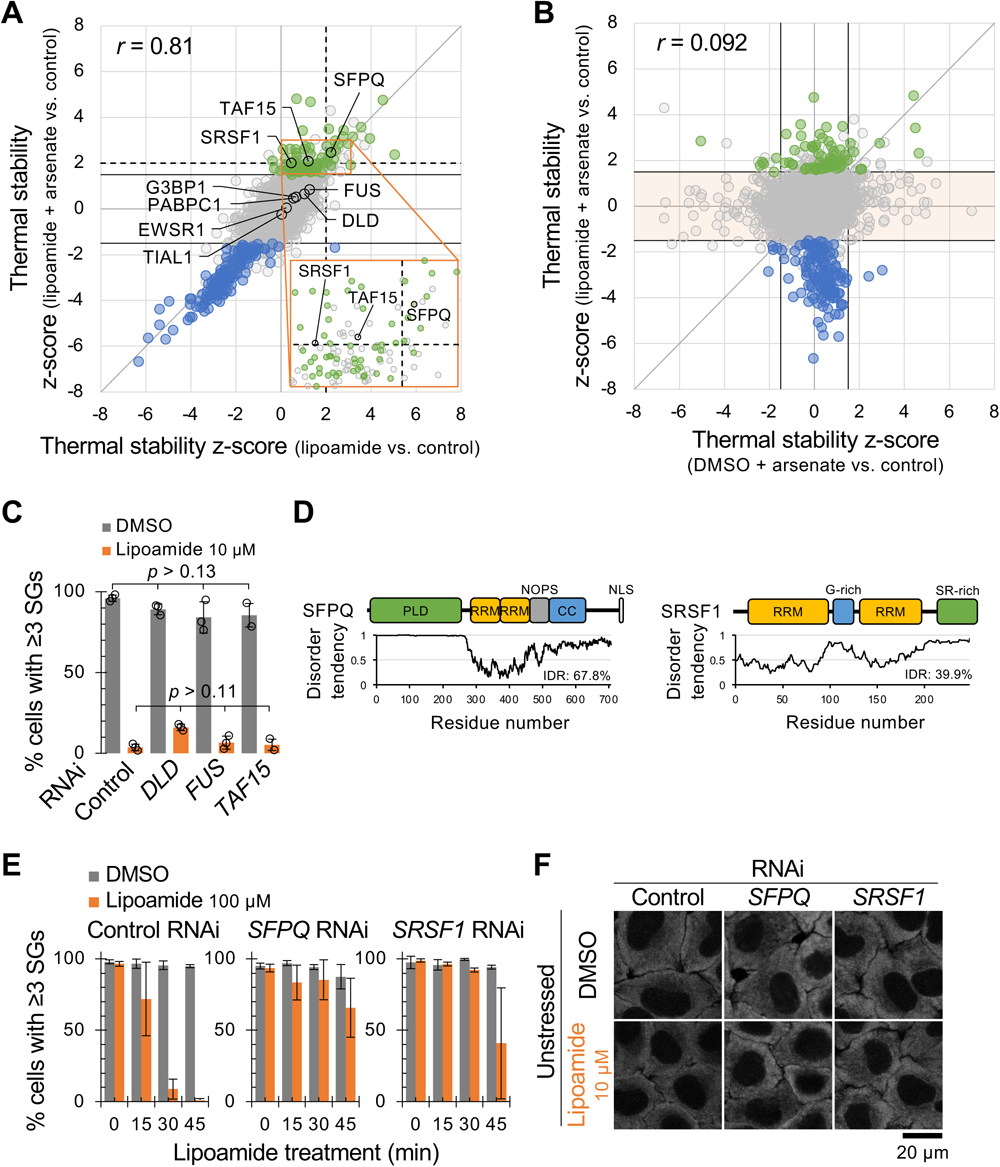
SFPQ and SRSF1 are cellular targets of lipoamide not necessary for stress granule formation. **A**, Z-scores of protein thermal stability in HeLa cells treated with only lipoamide and both lipoamide and arsenate. Proteins categorized as stabilized and destabilized in Fig. 4C,D are depicted in green and blue, respectively. The posìons of FUS, TAF15, DLD, SFPQ, and SRSF1 are indicated. Black solid and broken lines indicate cutoffs of z-scores used for the IDR analysis in Fig. 4C (±1.5) and the targeted RNAi screen (+2), respectively. **B**, Z-scores of protein thermal stability in HeLa cells treated with only arsenate and both lipoamide and arsenate. Black lines indicate |z-score| = 1.5. In most proteins with Increased or decreased thermal stability by only arsenate treatment (|z-score [arsenate]| > 1.5), the shifts were prevented by lipoamide pre-treatment (|z-score [arsenate + lipoamide]| < 1.5; masked in orange). Proteins categorized in stabilized and destabilized in Fig. 4C,D are depicted in the same colours; note that shifts in their thermal stability was not primarily due to treatment with arsenate. **C**, Mean ± s.d. of percentage of HeLa cells with ≥ 3 G3BP1-posìve stress granules. Cells depleted of indicated genes were treated with 10 µM lipoamide or 0.1% DMSO for 1 h followed by 1 mM arsenate for 1 h in the presence of lipoamide before stained with G3BP1. *n* = 324–393 cells from 3 independent experiments. *p* values by Tukey’s test. **D**, Domain composìons and distribùons of IDRs of human SFPQ (let) and SRSF1 (right). PLD, prion-like domain; RRM, RNA recognìon mòf; NOPS, NonA/paraspeckle domain; CC, coiled-coil domain; NLS, nuclear localizing signal; G-rich, glycine-enriched domain; SR-rich, serine/arginine-enriched domain. **E**, Mean ± s.d. of percentage of cells with ≥3 stress granules. HeLa cells depleted of indicated genes were treated with 3 mM arsenate for 1 h, and then with 100 µM lipoamide or the control DMSO in the presence of arsenate for indicated minutes. *n* = 213–467 cells from 3 independent experiments. **F**, Representàve images of HeLa cells treated and labeled as in Fig. 4E but without arsenate.

**Fig. S6.**
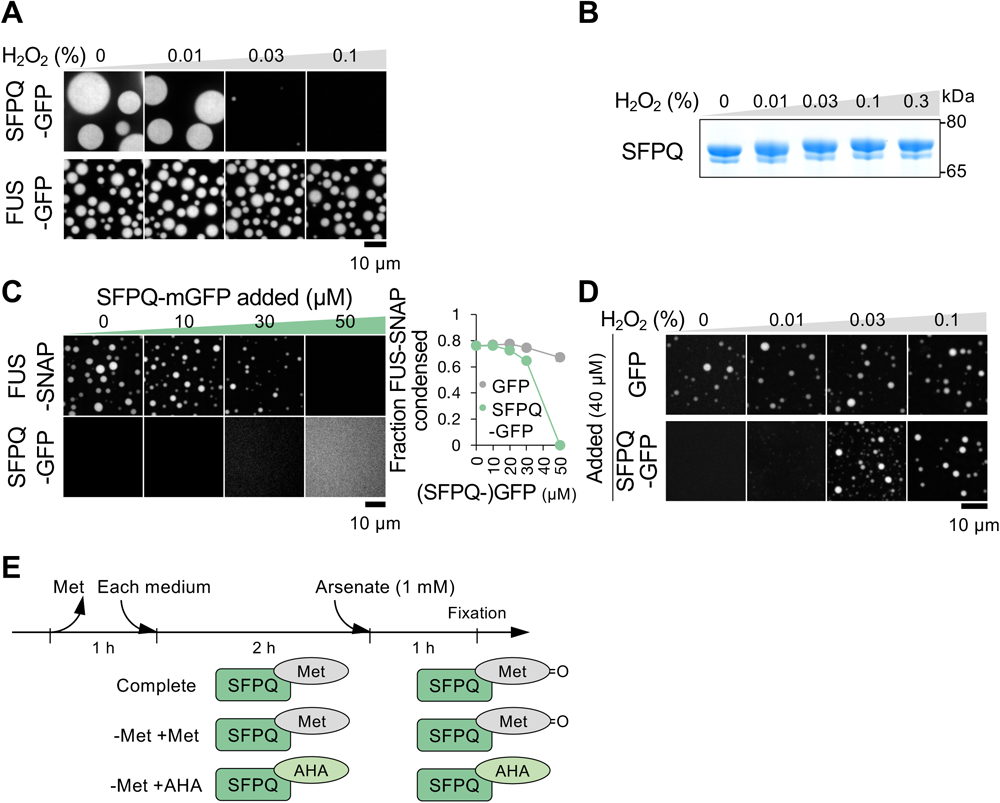
SFPQ dissolves FUS condensates *in vitro*. **A**, Representàve images of SFPQ-GFP (top, 10 µM) and FUS-GFP (bottom, 7 µM) protein condensates at a low salt condìon (75 mM KCl) and in the presence of H_2_O_2_ from >3 independent experiments. **B**, SDS-PAGE (non-reduced condìon) of 10 µM of the purified and untagged SFPQ proteins in diluted state oxidized with the indicated percentages of H_2_O_2_ for 30 min. **C**, Let, representàve images of co-incubàon of indicated concentràons of SFPQ-GFP and 6 µM of FUS-SNAP at a physiological salt concentràon (150 mM KCl) from >3 independent experiments. SFPQ proteins do not form condensates at 150 mM KCl while they suppress condensàon of FUS proteins in dosage-dependent manner. Right, mean ± s.d. of FUS condensate fraction in the presence of GFP (control) or SFPQ-GFP. *n* = 16 image fields. **D**, Representàve images of FUS-SNAP condensates (4 µM) co-incubated with 40 µM of GFP or SFPQ-GFP at a physiological salt concentràon (150 mM KCl) in the presence of indicated percentages of H_2_O_2_ from 3 independent experiments. **E**, Schema of the time course used in Fig. 5C. Cells were firstly cultured in methionine (Met)-free medium and then in each medium (complete medium or Met-free medium supplemented with Met or AHA) before arsenate treatment.

**Fig. S7.**
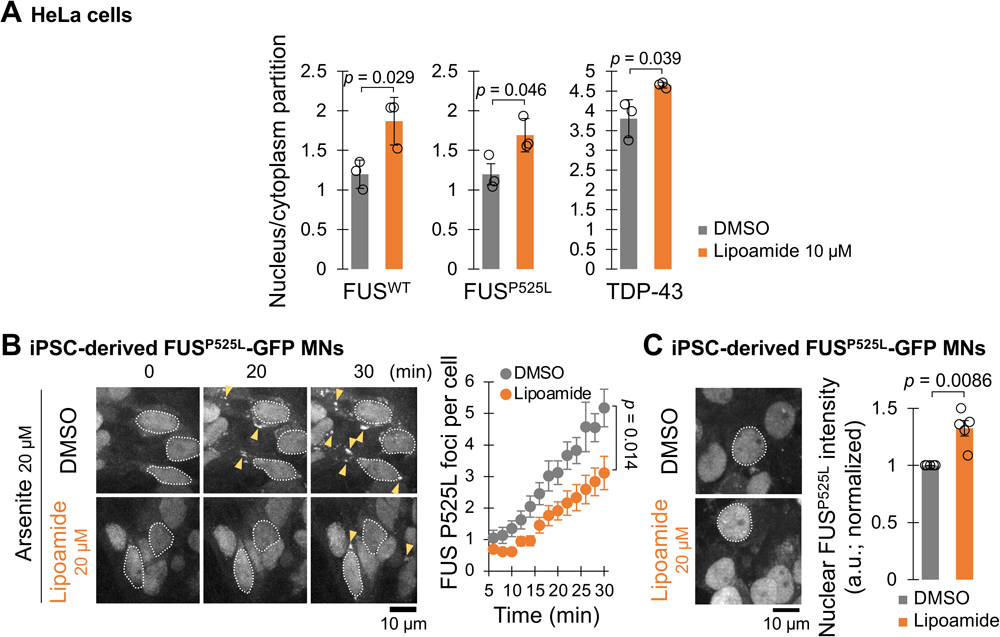
Lipoamide rescues nuclear localisation and functioning of ALS-linked proteins. **A**, Mean ± s.d. of nuclear-cytoplasmic intensity rào of FUS and TDP-43 in HeLa cells pre-treated with 10 µM lipoamide (0.1% DMSO as the control) for 1 h followed by 1 mM arsenate for 1 h in the presence of lipoamide. *n* = 290–603 cells from 3 independent experiments. *p* value by unpaired *t*-test. **B**, (Left) time-lapse images of iPSC-derived MNs expressing FUS P525L-GFP cultured for 14 days. Cells were treated with 0.02% DMSO or 20 µM lipoamide for 1 h followed by 20 µM arsenite for indicated minutes in the presence of lipoamide. Broken lines indicate outline of some nuclei. Arrowheads indicate some cytoplasmic FUS P525L foci. (Right) mean ± s.e.m. of number of FUS P525L foci per MN after arsenite treatment. *n* = 16 (DMSO) and 18 (lipoamide) cells from 3 independent experiments. *p* value by unpaired *t*-test. **C**, (Let) representàve images of iPSC-derived MNs expressing FUS P525L-GFP cultured for 5 days and then 30 days in the presence of 0.02% DMSO or 20 µM lipoamide. Broken lines indicate outline of some nuclei. (Right) mean ± s.e.m. of nuclear intensity of FUS P525L-GFP, normalized to that in the control (DMSO). *n* = 64–198 cells from 5 independent experiments. *p* value by one-sample *t*-test.

**Fig. S8.**
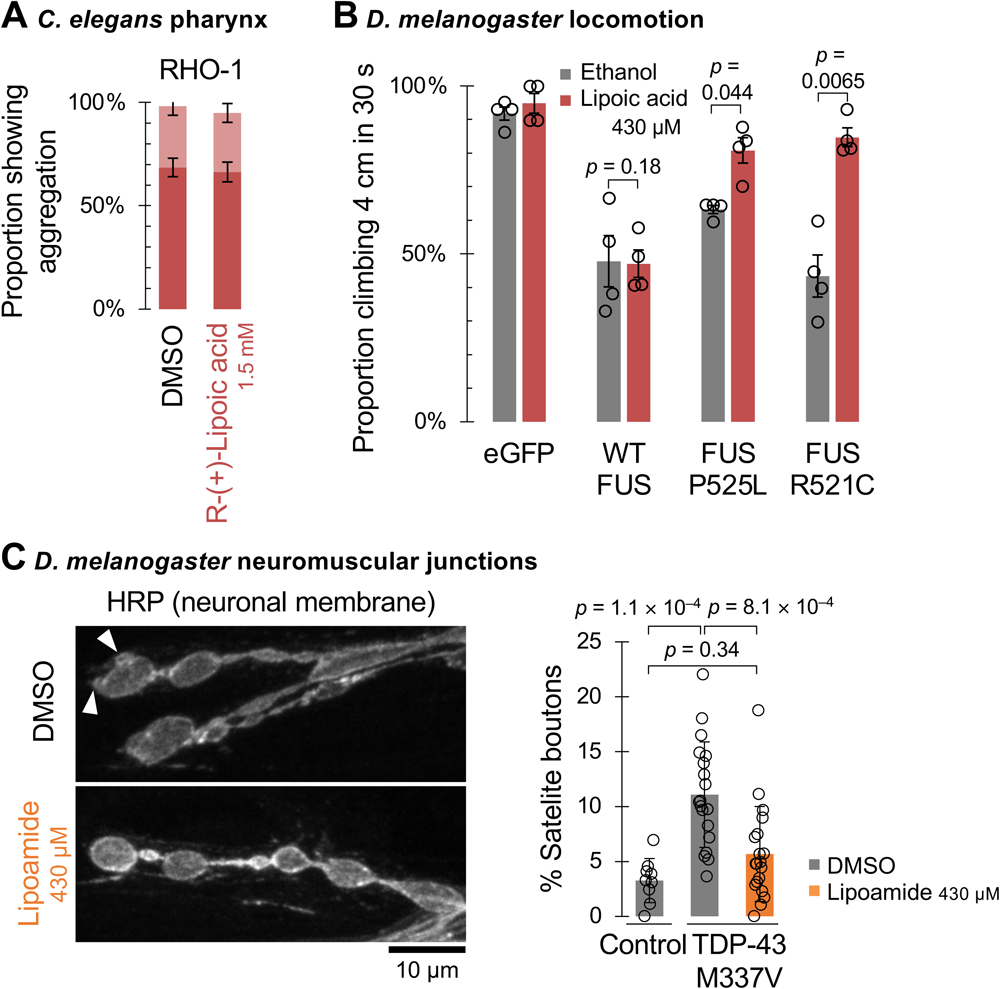
Extended analysis of *C. elegans* and *D. melanogaster* animal models of ALS. **A,** Mean ± s.e.m. of incidence of each protein aggregation in the pharyngeal muscles. Incidence of RHO-1 was scored on a low, medium, high scale (see methods). **B,** Mean ± s.e.m. of percentage of flies that climbed, with lipoic acid feeding in place of lipoamide in Fig. 6F. *p* values by unpaired *t*-test. **C,** (Left) Representative images of synaptic boutons of TDP-43 M337V-expressing flies, immunostained with an antibody against horseradish peroxidase (HRP), which labels the neuronal membrane. Arrowheads indicate appearance of satellite boutons. (Right) mean ± s.d. of percentage of satellite boutons (number of satellite boutons/number of total boutons) per fly. The control flies fed with 0.1% DMSO (grey; *n* = 9) and TDP-43 M337V-expressing flies fed with 0.1% DMSO (*n* = 19) or that containing 430 µM lipoamide (orange; *n* = 19) were examined. *p* value by Tukey’s test.

